# The Power of Three: Dynactin associates with three dyneins under load for greater force production

**DOI:** 10.1101/2025.01.14.632506

**Authors:** Lu Rao, Xinglei Liu, Mirjam Arnold, Richard J. McKenney, Kristy Stengel, Simone Sidoli, Florian Berger, Arne Gennerich

## Abstract

Cytoplasmic dynein is an essential microtubule motor protein that powers organelle transport and mitotic spindle assembly. Its activity depends on dynein-dynactin-cargo adaptor complexes, such as dynein-dynactin-BicD2 (DDB), which typically function with two dynein motors. We show that mechanical tension recruits a third dynein motor via an auxiliary BicD adaptor binding the light intermediate chain of the third dynein, stabilizing multi-dynein assemblies and enhancing force generation. Lis1 prevents dynein from transitioning into a force-limiting phi-like conformation, allowing single-dynein DDB to sustain forces up to ∼4.5 pN, whereas force generation often ends at ∼2.5 pN without Lis1. Complexes with two or three dyneins generate ∼7 pN and ∼9 pN, respectively, consistent with a staggered motor arrangement that enhances collective output. Under load, DDB primarily takes ∼8 nm steps, challenging existing dynein coordination models. These findings reveal adaptive mechanisms that enable robust intracellular transport under varying mechanical demands.

## INTRODUCION

Cytoplasmic dynein is the primary microtubule (MT) minus-end-directed motor in eukaryotic cells, playing essential roles in processes such as organelle transport, mitotic spindle positioning, and neuronal development^1–6^. Its activity is tightly regulated by cofactors, including lissencephaly-1 (Lis1), a protein critical for brain development^7–13^. Lis1 enables dynein to generate motion and force along MTs, and disturbances in Lis1 function—due to mutations or deletions— lead to type I lissencephaly^14,15^, a severe developmental brain disorder^16–18^. Despite its significance, the precise mechanisms by which Lis1 controls dynein function remain unclear, posing challenges for therapeutic advancements.

Dynein function is further modulated by interactions with dynactin, its largest co-factor. Dynactin not only enhances dynein’s processivity^1,4,19–29^—the ability to take consecutive steps without detaching—but also links dynein to cargoes and positions it within specific cellular domains^3,30,31^. Mutations in dynactin are associated with neurodegenerative diseases^32–34^ such as distal spinal bulbar muscular atrophy (dSBMA)^33^ and Perry syndrome^34^, highlighting its physiological importance. The coiled-coil cargo adaptor Bicaudal-D2 (BicD2) bridges the dynein tail and the dynactin shoulder^35,36^ (**Fig. 1A**), converting dynein from a diffusive^37^ or weakly processive^38,39^ entity into an ultraprocessive motor^24,25,40^. Recent studies have revealed that cargo adaptors like BicD2 can recruit two dynein motors to the dynein-dynactin-BicD (DDB-2) complex, forming a configuration of four parallel motor domains^41,42^ (**Fig. 1A**). This arrangement enhances force generation via load-sharing and improves processivity by reducing detachment from MTs^42^. The role of Lis1 within the DDB complex remains poorly understood. Initial studies suggested that Lis1 inhibits dynein motion by uncoupling ATPase activity from MT-binding dynamics^43–45^. More recent evidence indicates that Lis1 activates dynein by preventing its auto-inhibited “phi” state^46–53^, facilitating the recruitment of a second dynein dimer to the DDB complex^54,55^. Supporting this, Lis1 is thought to detach from dynein upon MT binding^56^, suggesting its primary role is motor recruitment rather than the regulation of active dynein motion. Despite recent structural and mechanistic insights into the function of dynein-dynactin-cargo adaptor complexes, key questions remain unresolved: How does force output scale with the number of dyneins in a complex? Studies of dynein-dynactin-BICDR1 (DDR) complexes have identified a second auxiliary BICDR1 molecule bound to dynactin as of yet unknown function. Furthermore, the interplay between Lis1-mediated dynein activation and motor recruitment under load is poorly understood, leaving significant gaps in our understanding of how these complexes dynamically adapt to varying intracellular force demands. Addressing these questions is essential to uncovering the principles governing dynein regulation and intracellular transport.

**Figure 1.**
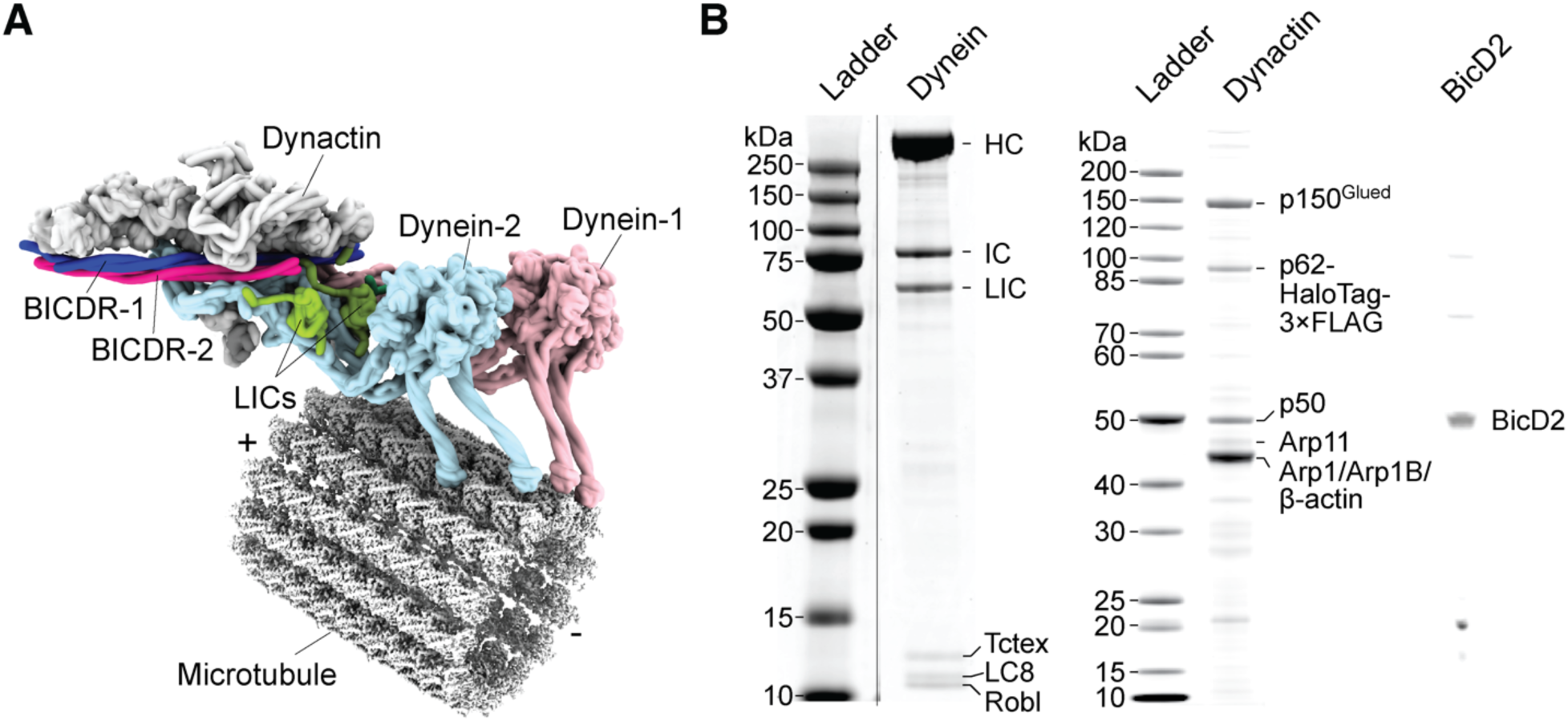
**(A)** Structural representation of dynein-dynactin-BicDR (DDR) complex (PDB 7Z8F^68^). The dynein heavy chain (B1) from the DDR complex was aligned with MTs (PDB 6DPV, EMDB 7974^101^), using the high-affinity dynein MT-binding domain structure bound to MTs (PDB 6KIQ^102^). **(B)** Polyacrylamide gel analysis of purified human dynactin complex, human dynein complexes, and BicD2(25-400). Protein bands were identified based on molecular weight, with the dynactin subunits further confirmed by mass spectrometry.

Here, we combined optical trapping, three-color single-molecule fluorescence, and structural analyses to uncover new principles governing dynein regulation within DDB complexes under mechanical tension. We reveal that Lis1 plays a dual role in dynein regulation: it recruits dynein motors to the DDB complex^54,55,57^ and activates single-dynein DDB (DDB-1) by prevention dynein from transitioning into a force-limiting “phi”-like conformation. This activation allows the motor to sustain forces up to ∼4.5 pN, compared to the ∼2.5 pN at which force generation typically terminates without Lis1. Notably, we discovered that DDB adapts to increasing load demands by recruiting a third dynein motor (DDB-3) via an auxiliary BicD adaptor that binds the light intermediate chain (LIC) of the third dynein. This second BicD molecule stabilizes multi-dynein assemblies, enabling higher force generation. The distinct forces generated by DDB-2 (∼7 pN) and DDB-3 (∼9 pN) suggest a staggered dynein arrangement, with load distributed unevenly among motors. Fully assembled DDB complexes primarily take ∼8 nm steps under tension, with smaller steps emerging near stalling forces. These findings provide mechanistic insights into Lis1-mediated dynein activation, the role of auxiliary adaptors in motor assembly, and the dynamic, tension-driven recruitment of dynein motors, offering a framework for understanding intracellular force generation and transport in complex cellular environments.

## RESULTS

### “Trains” of DDB Force Generation Reveal Distinct Stalling Forces

Previous studies on DDB force generation have primarily utilized non-native axonemes— 160-nm thick MT-containing structures purified from sea urchin^58^—and sub-stoichiometric dynein-dynactin-BicD concentrations (molar ratios of dynein-dynactin-BicD of 1:5:2^59^, 1:5:20^42,60^ and 1:2:10^54^). These conditions disfavored the formation of DDB complexes with two dyneins and limited the detection of distinct force-generation states.

For example, when DDB is tethered to trapping beads via a motor-less dynein tail, which permits only one dynein to bind while the second dynein-binding site is occupied by the tail fragment, the complex generates a force of 3.7 ± 0.2 pN (mean ± SEM) on axonemes^54^. This force matches that generated by DDB linked to beads via BicD^42^, suggesting that these studies primarily involved complexes with only a single dynein. Similarly, *S. cerevisiae* dynein alone generates 3.6 ± 0.2 pN on axonemes^59^ and 4.50 ± 0.04 pN on MTs^39,61^, indicating potential differences in DDB behavior on MTs.

To investigate the function of dynein-dynactin-adaptor (DDA) complexes with two dynein motors (DDA-2) (**Fig. 1A**), previous studies assembled motor complexes using the cargo adaptors HOOK3 and BICDR1, which are more likely than BicD to recruit two dyneins to dynactin^42^. As dynein-dynactin-HOOK3 (DDH, 4.9 ± 0.2 pN^42^) and dynein-dynactin-BICDR1 (DDR, 6.5 ± 0.3 pN^42^) generate more force than DDB (3.7 ± 0.2 pN^42^), it was reasoned that DDH and DDR recruit two dyneins. However, these studies were conducted on axonemes with a 1:5 molar dynein-dynactin ratio^42,60^, conditions under which, on average, only one in five dynactin complexes can bind a dynein. Thus, these complexes were unlikely to be fully saturated with two dyneins, leaving the source of their higher force generation unclear.

To address these limitations and gain insights into force production, we conducted optical trapping experiments using DDB complexes purified from rat brain lysate with a BicD-sfGFP fragment (amino acids 25–400)^24^ (**Fig. 2A**). Since neither dynein nor dynactin alone binds to BicD^24^, we anticipated that these complexes would contain dynein and dynactin in a 1:1 or 2:1 ratio. We then employed 500-nm trapping beads covalently linked to anti-GFP antibodies and used MTs instead of axonemes. Strikingly, we observed extended “trains” of force-generation events characterized by rapid force increases followed by stalling of movement (**Fig. 2C-F**). These force trains revealed four distinct stall forces for DDB: ∼2.5, ∼4.5, ∼7, and ∼9 pN (**Fig. 2B-F**). This discovery contrasts with earlier studies that reported broad detachment forces encompassing the plateaus we identified in our analyses (2–7 pN for DDB, 2.5–9 pN for DDH, and 3–11 pN for DDR)^51,55^. We speculate that the detection of these distinct plateaus in our experiments is due to the use of smaller beads, which, due to lower vertical forces, are likely to increase the DDB-MT on-rate and the likelihood of observing stalling (it’s worth noting that the effects of varying vertical forces on human dynein have not been tested, but higher vertical forces enhance MT detachment of kinesin-1 and prevent motor stalling^62,63^).

**Figure 2.**
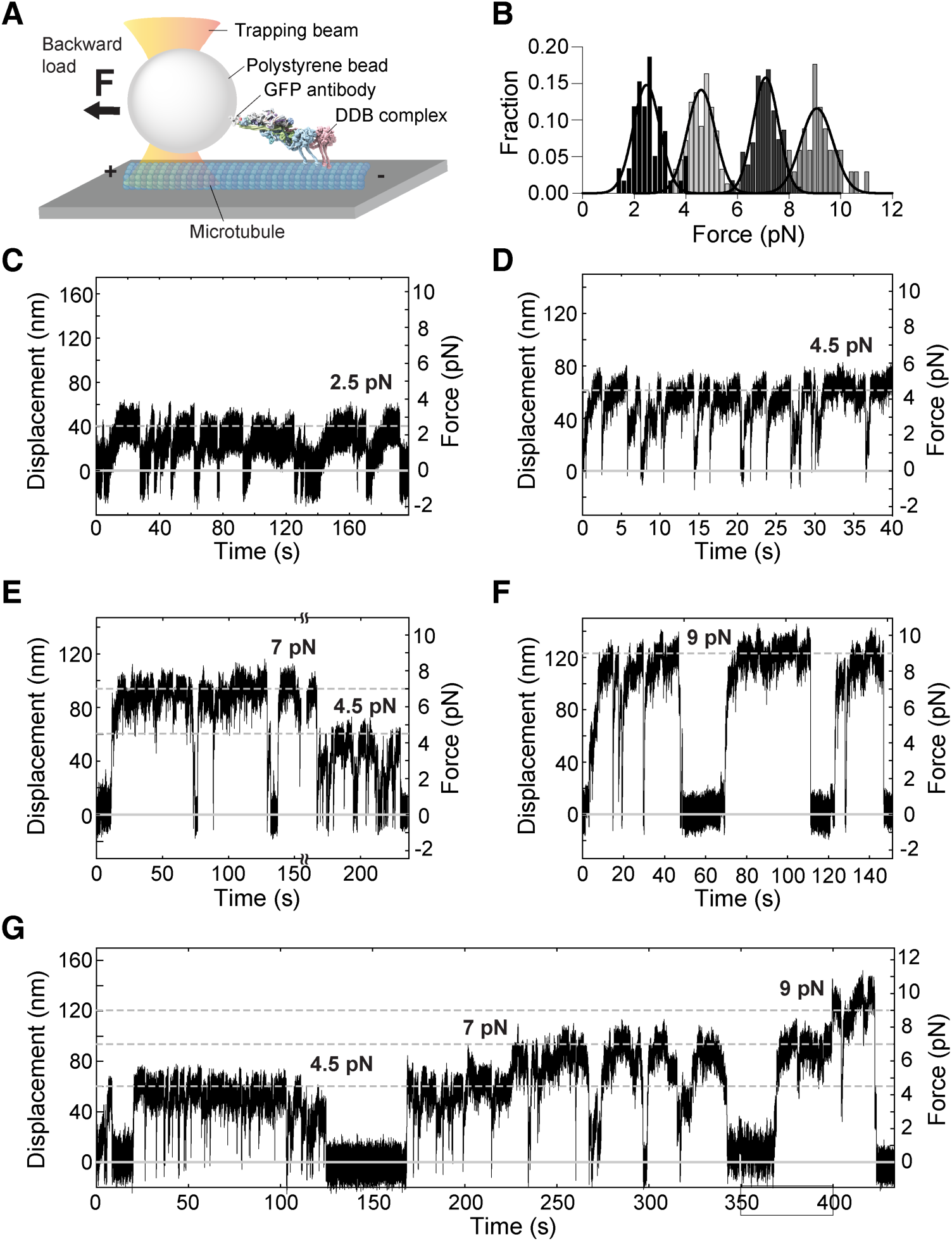
**(A)** Schematic of optical tweezers assay (not to scale). A ∼0.5 µm-diameter bead, coated with either anti-GFP antibody (as shown) or streptavidin, is tethered to DDB complexes via an sfGFP or biotin tag fused to the C-terminus of BicD2. The bead is positioned on a surface-attached microtubule. When the DDB binds to and moves along the MT in the presence of ATP, it pulls the attached bead with it. The optical trap resists this motion, generating a measurable force. **(B)** Stall forces of DDB complexes bound to anti-GFP antibody-coated beads. **(C–G)** Examples of DDB’s force generation.

### DDB-1 Stalls at ∼2.5 or ∼4.5 pN Depending on Its Partial or Full Activation

Since DDB complexes purified from mouse brain lysate demonstrated a myriad of generated forces (**Fig. 2C-F**), we sought to have a more controlled ratio of the assembling components to further investigate each force profile. Additionally, because we frequently observed only 1–2 events per bead for forces of ∼9 pN (**Fig. 2F**), we reasoned the binding strength between the anti-GFP antibody and GFP might not be strong enough to sustain large forces over time. To address this issue, we switched to using streptavidin-coated beads instead of anti-GFP-antibody-coated beads, taking advantage of the robust streptavidin-biotin linkage known to withstand forces up to 60 pN in force-extension experiments^64^.

We purified recombinant full-length human dynein, co-expressed with its five accessory chains, as previous desribed^25^. Biotinylated BicD (amino acids 25–400) was expressed in *E. coli* and purified (**Fig. 1B**). For dynactin, traditionally purified from bovine, porcine, rodent brain tissue^19,24,25^, we employed a CRISPR-based approach to tag the DCTN4 subunit with a HaloTag-3×FLAG at its C-terminus in a human suspension culture cell line (HEK293sus), following established protocols^65^. This tagging strategy, recently used to generate a stable adherent HEK293 cell line^66^, enabled efficient recombinant expression of dynactin (**Fig. 1B**). With each component—dynein, BicD2, and dynactin—expressed and purified to high purity and concentration, we successfully assembled DDB complexes in defined stoichiometric ratios.

To investigate the force generation of DDB when associated with a single dynein, we assembled the complexes in a 1:1:1 ratio of dynein, dynactin, and BicD. Our previous studies have indicated that isolated human dynein can transition from slow movement toward ∼2.5 pN to a rapid movement toward ∼4 pN (**Fig. 3A**, reproduced from Fig. S3 in ref^39^). We observed a similar behavior in DDB complexes, transitioning from slow movement toward ∼2.5 pN followed to rapid movement toward ∼4.5 pN (**Fig. 3B**). Interestingly, for human dynein alone, we only observed instances of transitioning from ∼2.5 to ∼4 pN under increased trap stiffnesses (∼0.06 pN/nm); at a lower trap stiffness of 0.01 pN/nm, it stalls at ∼1 pN (**Fig. 3A**, inset, and Fig. S3 in ref^39^), suggesting that a higher force per unit displacement facilitates the full activation of dynein. This indicates that the two distinct forces are generated by a single dynein altering its behavior depending on its activation state.

**Figure 3.**
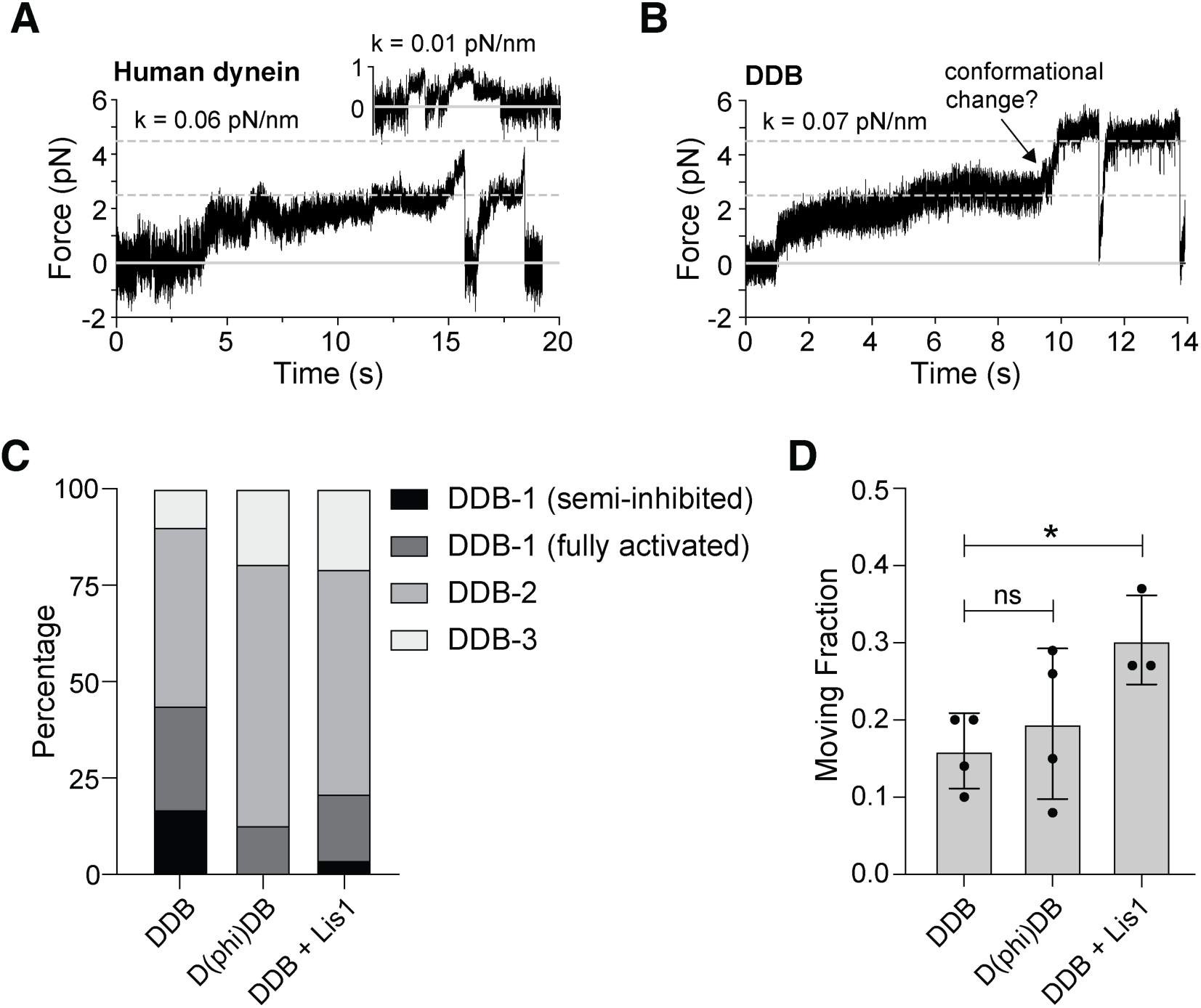
**(A)** Force generation of human dynein in a stiff optical trap (adapted from^39^). **(B)** Example of force generation by DDB-1, showing a transition from slow movement toward ∼2.5 pN to rapid movement toward ∼5 pN. **(C)** Distribution of stall forces (∼2.5, ∼4.5, ∼7 and ∼9 pN) across different experiments: DDB-1(semi-inhibited), DDB-1 (fully activated), DDB-2, and DDB-3. The percentages represent the fraction of beads exhibiting each stall force, with sample sizes and distributions as follows: DDB: n=41; 17%, 27%, 46%, 10%. D(phi)DB: n=31, 0%, 13%, 68%, 19%. DDB+Lis1: n=29; 3%, 17%, 59%, 21%. **(D)** The measured fraction of force-generating beads. The bars represent the mean value with SD. DDB: n=4, 0.16 ± 0.05 (mean ± SD). D(phi)DB: n=4, 0.19 ± 0.10; DDB + Lis1: n=3, 0.30 ± 0.06.

Recent studies have shown that mammalian dynein often adopts an auto-inhibited conformation, commonly referred to as the “phi” particle^48–52^, which hinders efficient interactions with MTs. Unlike mammalian dynein, *S. cerevisiae* dynein moves processively along MTs by itself^67^ and a mutant form unable to assume the phi conformation exhibits an increased run length along MTs^46^, indicating that the auto-inhibited state limits the movement of wild-type *S. cerevisiae* dynein on MTs. To investigate whether a similar auto-inhibited state restricts in human dynein’s force generation to ∼2.5 pN, we assembled DDB complexes at a 1:1:1 ratio with a dynein mutant (K1610E and R1567E double mutant^48^) incapable of adopting this inhibitory conformation, referred to here as D(phi)DB. Contrary to expectations, D(phi)DB complexes predominantly stalled at ∼7 pN (**Fig. 3C** and **Suppl. Fig. 1A**), even when assembled at a molar ratio of 1:6:6 or in the presence of excess dynein tail (**Suppl. Fig. 1B&C**), suggesting that dynein in its open conformation tends to cluster and form DDB complexes with two dyneins per complex (discussed further in the following section). However, this dynein mutant prevents stalling at ∼2.5 pN in the lower force regime, with only ∼4.5 pN forces observed (**Fig. 3C** and **Suppl. Fig. 1A&B**). Moreover, the regulatory protein Lis1, which prevents dynein’s auto-inhibited state^46,47,53–55^, significantly reduces the occurrence of stalling at ∼2.5 pN (**Fig. 3C**). These results confirm that a DDB complex bound to a single dynein stalls at either ∼2.5 or ∼4.5 pN depending on whether it is partially or fully activated.

### Fully active DDB-2 stalls 7 pN

D(phi)DB complexes generate forces of ∼4.5, ∼7, or ∼9 pN (**Suppl. Fig. 1A**), suggesting that fully active DDB-1 generates ∼4.5 pN, while a DDB complex with two dyneins (DDB-2) generates ∼7 pN. This observation aligns with that of DDR, which binds two dyneins preferentially and is reported to generate 6.5 pN^42^.

A key question is what accounts for the observed stalling at 9 pN. If fully active DDB-1 generates ∼4.5 pN, then logically, fully active DDB-2 might be expected to reach up to ∼9 pN if both motors contribute equally to force generation. However, DDR assembled with the dynein mutant and in the presence of Lis1—both intended to prevent the phi conformation—has been observed to generate only 6.1 pN^54^, which aligns more closely with the ∼7 pN force we measure. This discrepancy suggests that the ∼7 pN stall force does not arise from a collaboration between a fully active dynein and a partially active motor. Instead, it likely results from uneven load distribution between the motors, with the leading motor bearing a greater load than the lagging dynein, contributing only ∼2.5 pN.

To further support the hypothesis that the ∼7 pN force is generated by two actively engaged dyneins, we conducted experiments by introducing 5 nM dynein into a slide chamber containing pre-assembled 1:1:1 ratio DDB complexes bound to trapping beads. This resulted in ∼60% of the moving beads displaying forces of ∼7 pN (**Fig. 4E** and **Suppl. Fig. 2**). Notably, the proportion of beads showing force generation remained consistent with that observed in DDB-only complexes (∼10%), despite the addition of free dynein (**Fig. 4D**). This indicates that the freely diffusing dyneins selectively bind to DDB complexes anchored via the biotinylated BicD to the bead surfaces, rather than non-specifically binding to the bead surfaces, which would have increased the fraction of force-generating beads^39^.

**Figure 4.**
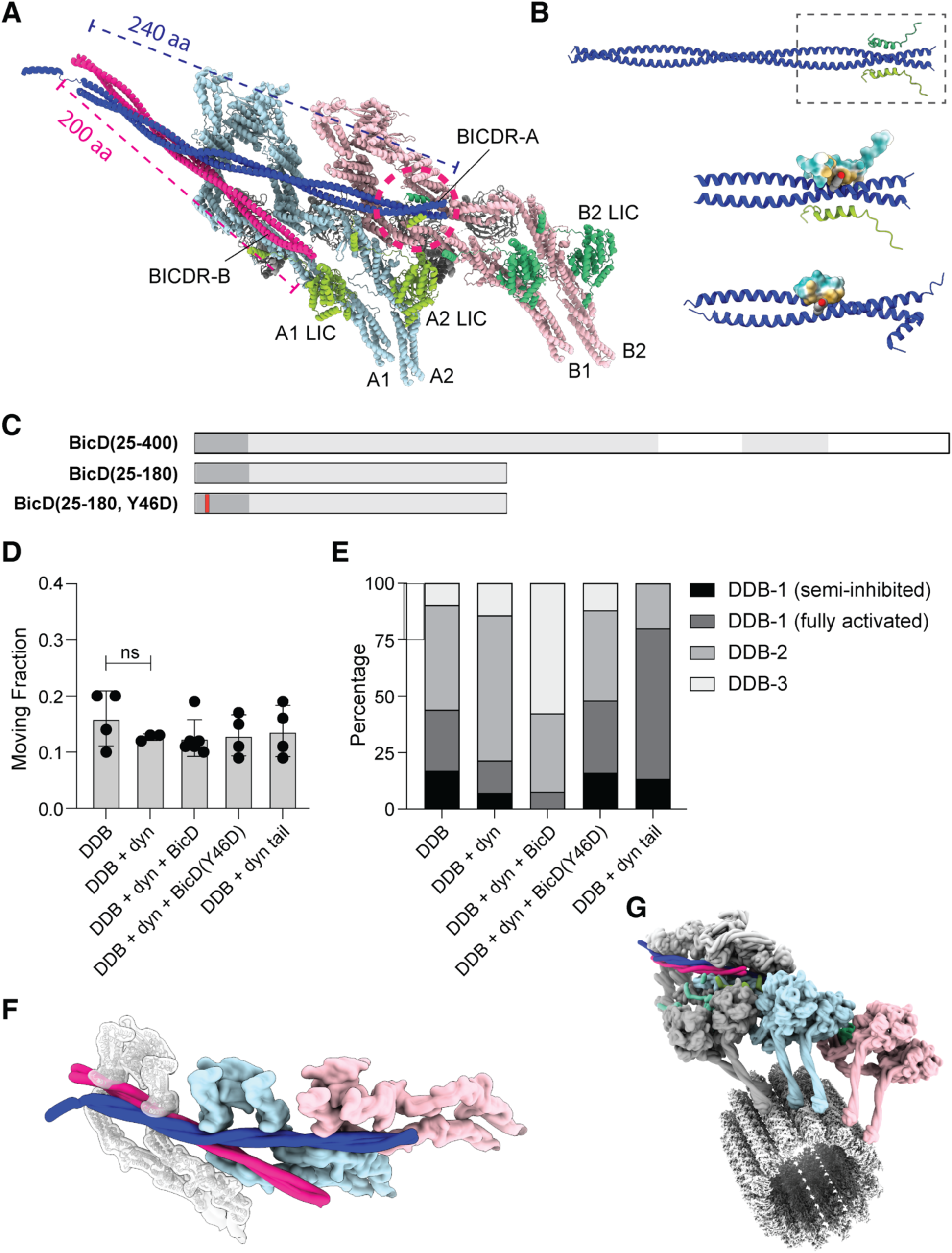
**(A)** Structure of DDR complex with two dyneins (light blue and pink) and two BicDR dimers (blue and magenta) (PDB 7Z8F). Dynactin is not shown in the representation. The interaction between the C-terminal helix of the LICs of dynein A2 (light green) and B1 (green) with the N-terminal CC1-box of BicDR-A (blue) highlighted with the dashed red circle. The span of BicDR across the dynactin shoulder is indicated with corresponding colors. **(B)** (Top) ColabFold-predicted^99^ structure of the interactions between mouse BicD2(25–180) and human dynein LIC(417–444). (Middle) A close-up of the predicted structure, showing Tyr46 of one BicD2 as sphere style (colored by elements). The hydrophobic surface of one LIC is depicted in light yellow (hydrophobic) and cyan (hydrophilic). (Bottom) Crystal structure of human BicD2(1– 98) binding to human dynein LIC(433–458), displayed with hydrophobic surface (PDB 6PSE^70^). **(C)** Schematic representation of the BicD constructs. Y46D mutation in BicD2(25–180) is marked in red. **(D)** The moving fraction of the beads per experiment. The bars represent the mean value with SD. DDB: n=4, 0.16 ± 0.05 (mean ± SD). DDB+dyn: n=3, 0.13 ± 0.01. DDB+dyn+BicD: n=6, 0.12 ± 0.03. DDB+dyn+BicD(Y46D): n=4, 0.13 ± 0.04. DDB+dyn tail: n=4, 0.14 ± 0.05. **(E)** The percentage for each DDB complex. The percentages represent DDB-1(semi-inhibited), DDB-1 (fully activated), DDB-2, and DDB-3, respectively. DDB: n=41; 17%, 27%, 46%, 10%. DDB+dyn: n=14, 7%, 14%, 64%, 15%. DDB+dyn+BicD: n=26, 0%, 8%, 35%, 57%. DDB+dyn+BicD(Y46D): n=25, 16%, 32%, 40%, 12%. DDB+dyn tail: n=15, 13%, 67%, 20%, 0%. **(F)** Hypothetical structure of a third dynein dimer (gray, semi-transparent) recruited to a DDB complex. The tail was manually added to the DDR structure. **(G)** Full DDR complex, including a microtubule, shown for context.

### Auxiliary BicD Enables Third Dynein Recruitment and Enhanced Force Generation

Given the observed force trains at ∼9 pN and the likely contribution of two fully active dyneins to the ∼7 pN force, we hypothesized that the ∼9 pN force results from the functional recruitment of a third dynein to the DDB complex. Although this possibility has not been previously explored, the cryo-EM structure of two dyneins bound to dynactin and BICDR1^42^ suggests that the dynactin shoulder provides sufficient space for a third dynein tail to bind (**Fig. 4F&G**).

To test this hypothesis, we examined whether dynactin can accommodate a third dynein by generating two truncated BicD constructs, BicD(25–276) and BicD(25–180), based on the cryo-EM structure of DDB^42^. BicD(25–276) spans the full shoulder of dynactin, encompassing the putative third dynein-binding site (**Fig. 4A&F**), whereas BicD(25–180) extends only to the position occupied by the middle dynein (dynein-A^42^) (**Fig. 4A&F**). Using single-molecule fluorescence and optical-tweezers assays, we assessed the functionality of these constructs. BicD(25–276) formed DDB complexes capable of motion and force generation (**Suppl. Fig. 3A**), similar to BicD(25–400) (**Fig. 2**). In contrast, the shortest construct, BicD(25–180), not only failed to form DDB complexes with two dyneins (DDB-2) but also formed significantly fewer complexes with one dynein (DDB-1) (**Suppl. Fig. 4**), generating forces of only ∼2.5 or ∼4.5 pN (**Suppl. Fig. 3B**). These results indicate that BicD’s ability to span the full dynactin shoulder is essential for forming and stabilizing DDB complexes, although the terminal segment (amino acids 277–400) does not interact with dynein directly. Consequently, the essential role of the middle segment (amino acids 180–276) leaves open the question of whether it participates in the binding of the third dynein.

A recent cryo-EM study revealed that a second BicDR1 molecule (BICDR-B) can stabilize the MT-bound DDR complex by acting as an auxiliary adaptor^68^ (**Fig. 4A**). While a second BicD molecule has not yet been visualized in DDB cryoEM structures, we investigated the possibility that a second auxiliary BicD adaptor facilitates the recruitment of the third dynein by employing the shortest construct, BicD(25–180), without a biotinylated C-terminus. This modified construct, incapable of binding to streptavidin beads or forming functional DDB complexes on its own, retains the segment capable of interacting with pre-assembled DDB complexes that include BicD(25–400) anchored to streptavidin beads. When 5 nM free dynein and 5 nM BicD(25–180) were added to DDB complexes assembled at a 1:1:1 ratio, we observed dominant ∼9 pN forces in ∼60% of all force-generating beads (**Fig. 4E** and **Suppl. Fig. 5**). These results strongly suggest that a second auxiliary BicD adaptor plays a critical role in recruiting a third dynein.

Further insights into this interaction came from cryo-EM structures of DDR complexes with two BicDR1 molecules^69^ (**Fig. 4A**). The CC1-Box (AAxxG; “x” marks any amino acid) of the N-terminus of the primary BicDR1 (BICDR-A) interacts with α-helix-1 of the C-terminus of the light intermediate chain (LIC) from dynein B (B1)^69^ and with the LIC of dynein-A (A2) (**Fig. 4A&B**). Indeed, removing the N-terminus of BicD completely abolished the formation of DDB complex (**Suppl. Fig. 4**). In contrast, the auxiliary BicDR1 (BICDR-B) N-terminus interacts exclusively with the LIC of dynein-A2^69^. Although the LIC of dynein-A2 interacts with both BICDR-A and BICDR-B^69^, the latter’s unoccupied surface may bind the LIC of the third dynein. To directly assess whether disrupting the interaction between the auxiliary BicD and the third dynein’s LIC impairs recruitment, we introduced a single-point mutation (Y46D) in BicD(25– 180). This mutation disrupts the hydrophobic interaction between the BicD N-terminus and the dynein LIC^70^. When 5 nM mutated BicD(25–180, Y46D) and 5 nM free dynein were added to pre-assembled DDB complexes, we observed a significant reduction in force generation, with less than 13% of beads producing ∼9 pN forces (**Fig. 4E**). These findings demonstrate that recruitment of the third dynein critically depends on the interaction between the second auxiliary BicD and the third dynein’s LIC.

### Tension-Driven Recruitment of a Third Dynein to DDB Complexes

A potential question arises: why have cryo-EM and cryo-ET studies not observed the presence of a third dynein, despite its capability to associate with DDB? The explanation likely lies in the experimental conditions. Unlike structural studies, our optical-trapping assay applied mechanical load to DDB complexes (**Fig. 2A**), and this mechanical tension revealed a step-like increase in stalling forces (**Fig. 2G**). This intriguing phenomenon suggests that mechanical tension facilitates the recruitment of additional dynein motors to DDB complexes, enabling adaptation to increased force demands.

To investigate this concept, we engineered a kinesin-DDB chimera by covalently linking a DDB complex through BicD to the C-terminal tail domain of a kinesin-1 rigor mutant^71^ using the SpyTag-SpyCatcher system^72^. Specifically, we inserted the SpyCatcher (SpyCatcher003^73^) at the C-terminus of BicD—typically bound to cargo—and fused the SpyTag (SpyTag003^73^) to the C-terminus of the kinesin mutant (**Fig. 5A&B**). This rigor kinesin mutant is characterized by its strong binding to MTs and lack of motility^71^. We hypothesized that if the kinesin-DDB chimera attaches to the MT with the kinesin motor domains and the DDB complex bound to the MT, the DDB complex must generate enough force to detach the kinesin motor domains from the MT, allowing it to move toward the MT minus-end. Once the generated force reaches a certain threshold, a third dynein could potentially bind to the DDB complex (**Fig. 5A**).

**Figure 5.**
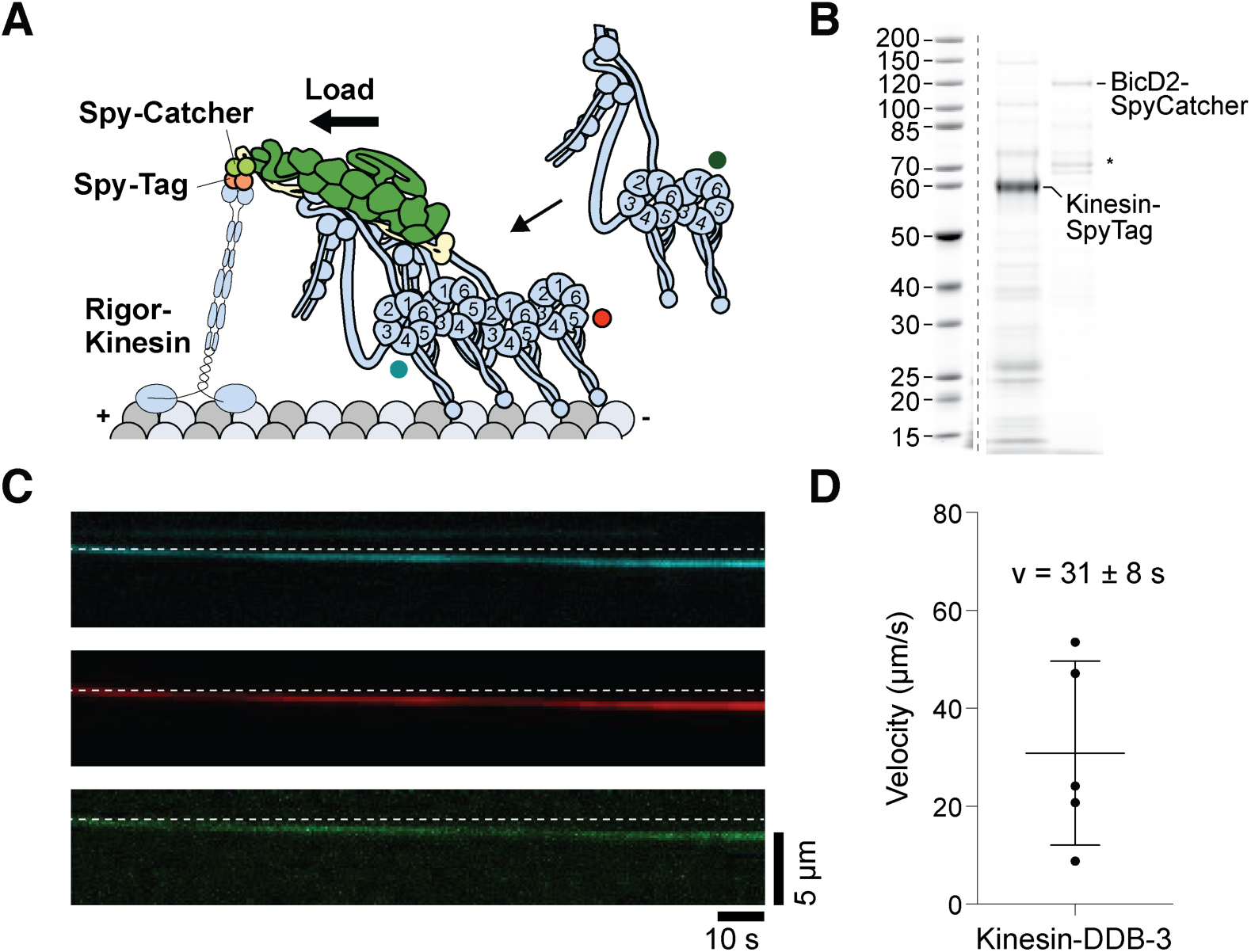
**(A)** Schematic of a rigor kinesin-1 covalently attached to BicD in the DDB complex via SpyTag-SpyCatcher system. **(B)** Polyacrylamide gel analysis of the rigor kinesin–BicD construct. (*) denote side products with truncated kinesin tails, which do not interfere with the experiments. **(C)** Representative kymograph of a kinesin-DDB complex containing three different dye-labeled dynein moelcules. **(D)** The velocity of the three-color colocalized complexes at 100 µM ATP. The bars indicate mean with SD (0.031 ± 0.018 µm/s).

To test this hypothesis, we pre-assembled DDB complexes with two differently labeled dyneins (Alexa Fluor 647 and TMR dyes) and introduced these dual-labeled DDB complexes—expected to display two different colors in 50% of cases—into the slide chamber with free dynein labeled with a third color (Alexa Fluor 488). Using three-color, single-molecule total internal reflection fluorescence (smTIRF) microscopy, we observed that moving chimeras were frequently associated with three dyneins (**Fig. 5C**). In contrast, DDB complexes lacking the kinesin did not recruit a third dynein. The average velocity of ∼30 nm/s (**Fig. 5D**) aligns with the movement of a DDB-3 complex under a load above ∼8 pN (**Suppl. Fig. 6**), further supporting the role of tension in motor recruitment.

In conclusion, our findings demonstrate for the first time that mechanical tension applied to a DDB complex facilitates the recruitment of a third dynein. This load-induced adaptation leads to augmented force generation and increased velocities, enabling the DDB complex to meet escalating mechanical demands.

### DDB takes largely load-independent steps of ∼8 nm along MTs

Cryo-EM studies suggest that the motor domains of the two dyneins engaged within DDB complexes adopt a relatively compact and parallel arrangement^41^, which would predict small center-of-mass steps along MTs. In contrast, previous single-molecule fluorescence studies have reported highly variable step sizes for DDB complexes in the absence of load, ranging from 4 to 48 nm^54,60^. Optical trapping experiments further revealed that the average forward step size of DDB complexes decreases with increasing load, from ∼15 nm (step sizes ranging from 4 to 32 nm) at a 0.4 pN load to ∼10 nm (step sizes ranging from 2 to 28 nm) at a 3.6 pN load^60^. Given the structural arrangement of the dynein motor domains^41^, steps sizes as large 48 nm would require substantial rearrangements of the motor domains, making such displacements unlikely.

In our experiments, DDB complexes bound to a single dynein predominantly advanced in ∼8-nm center-of-mass steps under loads exceeding ∼1 pN, even after accounting for the complex’s compliance (**Suppl. Fig. 7**). At lower loads, consecutive forward steps occasionally extended to ∼16 nm (**Fig. 6B**, right top, red). For DDB complexes associated with two dyneins, the step sizes remained largely consistent at ∼8 nm under loads above ∼2 pN, with occasional steps reaching up to ∼24 nm under low-load conditions (**Fig. 6B**, right middle, red). These findings align with previous step-size measurements in the absence of load^54,60^. In contrast, when DDB complexes were associated with three dyneins, the step sizes were smaller than those of DDB-1 and DDB-2 (**Fig. 6B**, right bottom, red), suggesting that the third dynein influences the movements of the two leading motors, thereby reducing the center-of-mass displacements.

**Figure 6.**
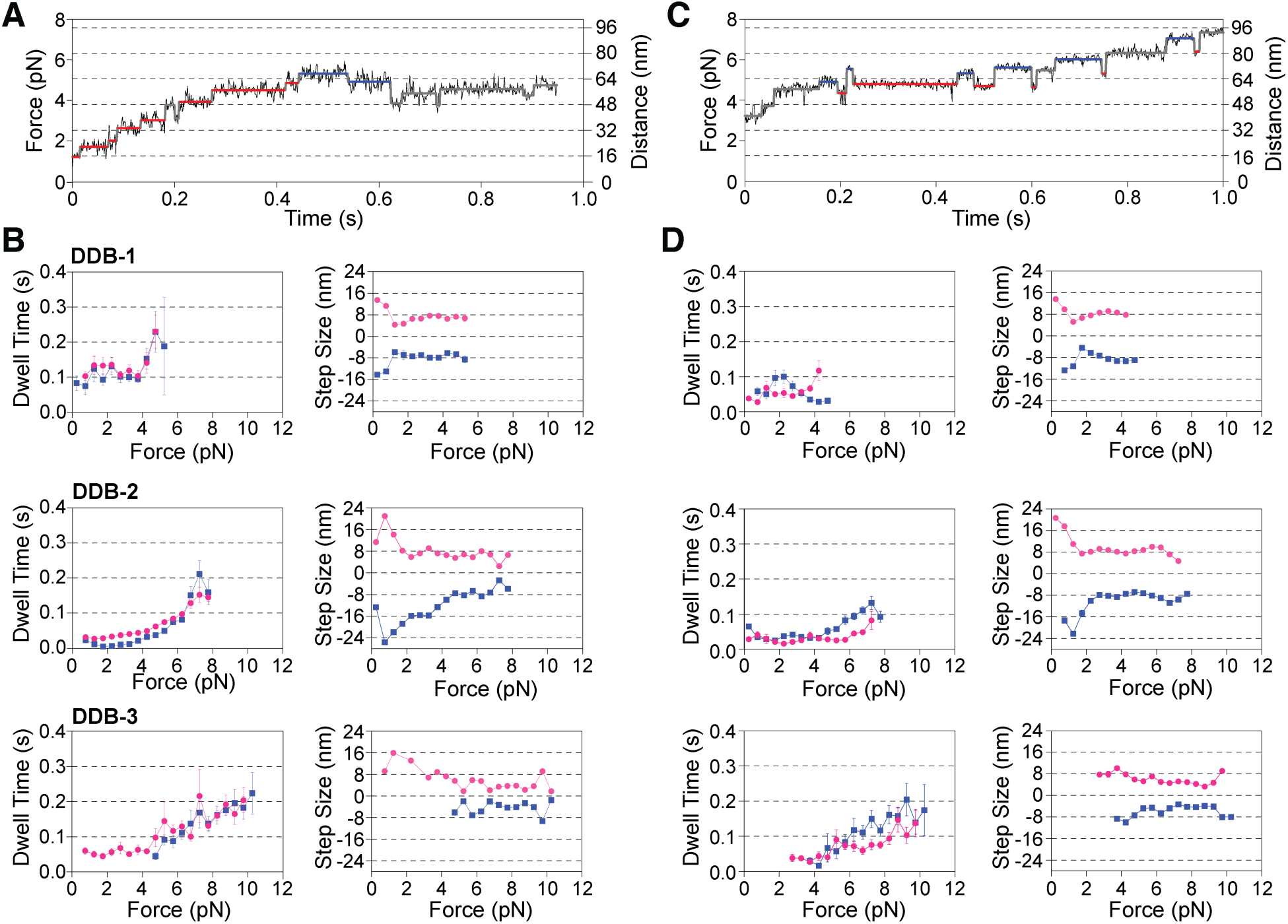
**(A)** Example trace of DDB-2 taking consecutive steps. Red segments indicate the dwell time before consecutive forwards steps, while blue segments indicate the dwell time before consecutive backward steps. **(B)** Relationship between dwell time, step sizes of consecutive steps, and applied forces for DDB complexes. Forward steps are shown in red, and backward steps in blue. Data were binned using a 0.5-pN force window. Bars represent the mean ± SEM. **(C)** Example trace of DDB-2 taking oscillatory steps. Red segments indicate the dwell time before forward steps, while blue segments indicate the dwell time before backward steps. **(D)** Relationship between the dwell time, step sizes of oscillatory steps, and applied forces for DDB complexes. Forward steps are shown in red, and backward steps in blue. Data were binned using a 0.5-pN force window. Bars represent the mean ± SEM.

For consecutive backward steps (**Fig. 6B**, right, blue), DDB-1’s exhibited a backward stepping behavior similar to its forward stepping (**Fig. 6B**, right top, blue). In contrast, DDB-2 displayed larger backward step sizes under loads below 4 pN (**Fig. 6B**, right middle, blue). This observation suggests that while DDB-1’s backward step size may be constrained by the load-induced geometry of its two motor domains, the two dynein dimers of DDB-2 can separate by greater distance at low load. Such large backward movements may facilitate synchronization between the dynein dimers if the leading dimer advances too far ahead. Conversely, DDB-3 showed almost no backward stepping under low-load conditions (**Fig. 6B**, right bottom, blue), likely because, in this configuration, at least two dynein dimers remain bound to the MT at a given time, minimizing backward stepping.

In addition to the forward-advancing steps and subsequent backward steps described above (**Fig. 6A**), we also observed an “oscillating” forward-backward stepping behavior reminiscent of the load-induced stepping behavior of *S. cerevisiae* dynein^74^ (**Fig. 6C**). Both stepping behaviors were observed across all DDB complexes. While the step sizes in the two stepping modes are comparable (**Fig. 6B & D**, right), their dwell time profiles differ significantly (**Fig. 6B & D**, left). In the oscillating stepping mode, a backward step is often immediately followed by a forward step (**Fig. 6C**), with shorter dwell times preceding the subsequent forward step and reduced force dependency compared to advancing stepping. This suggest that, unlike kinesin-1, the stepping order of DDB’s motor domains is more stochastic with similarities to the stepping behavior of *S. cerevisiae* dynein^67,74–76^.

Together, our step-size analysis reveals that DDB complexes predominantly advance in ∼8-nm center-of-mass displacements under sub-stall loads, reflecting a compact and efficient motor arrangement. Under near-unloaded conditions, DDB complexes exhibit their full range of step sizes, extending up to ∼24 nm. Conversely, as loads approach stalling, steps become smaller, occasionally shrinking to ∼4 nm, suggesting an inching mechanism that allows the complex to adapt to mechanical resistance with minimal displacement increments. These findings indicate that the motor domains within DDB complexes maintain a tightly coordinated, compact configuration under load, while low-load conditions allow for limited splaying or flexibility. This dynamic behavior underscores the capacity of these complexes to adjust their stepping mechanics in response to varying mechanical demands.

## Discussion

Our study reveals a remarkable mechanosensitive property of DDB motor complexes: under increasing load, additional dyneins can be recruited, enhancing both force production and processivity. By integrating recombinant protein expression, structural analyses, optical trapping, and multi-color single-molecule fluorescence imaging, we demonstrate how mechanical tension triggers the higher-order assembly of DDB complexes. These findings expand our understanding of how dynein adapts to the mechanical demands of intracellular transport and unveil novel mechanisms of load-dependent motor recruitment and coordination.

### Dynein Force Generation Is Limited by Auto-Inhibition

For nearly two decades, optical trapping experiments have indicated that mammalian dynein, when isolated from cofactors, produces forces of ∼1 pN or less when measured with a weak optical trap (stiffness ∼0.01 pN/nm)^39,52,77^. Forces of ∼7–8 pN have been reported^78,79^, but the basis for these outliers remain unclear. At higher trap stiffness (0.1 pN/nm) —conditions where dynein’s weak processivity has less impact—its force increases to ∼2 pN^39^, consistent with a link between its weak processivity and limited force generation^39^. By contrast, *S. cerevisiae* dynein exhibits high processivity and a trap-stiffness-independent stall force of 4.5 pN^39^.

Our data reveal that mammalian DDB complexes with a single dynein (DDB-1) generate forces of ∼2.5 pN when dynein transitions into a conformation that ends a run (at a trap stiffness of 0.06 pN/nm). This value closely aligns with the ∼2 pN stall force reported for isolated dynein^39,59^, suggesting that dynein’s limited force generation stems from its tendency to adopt an auto-inhibitory “phi”-like conformation. In this state, the motor stops advancing under loads of ∼2.5 pN, precluding higher force output. Although auto-inhibited dynein has a significantly reduced MT-binding affinity^48^, it is plausible that when one of the two motor domains is MT-bound and bearing load, the detached motor domain attempts to adopt the phi-like conformation by crossing its stalk over the stalk of the bound domain. If the primary AAA+ ATPase domain (AAA1) is nucleotide-free, the MT-bound motor domain may form a force-insensitive ideal bond, as demonstrated for *S. cerevisiae* dynein^61,80,81^. Notably, DDB-1 complexes carrying mutations that disrupt auto-inhibition exclusively generate forces of ∼4.5 pN. This strongly supports the idea that auto-inhibition represents the primary limitation on the force generation of a single dynein.

Lis1 acts as a critical regulator by counteracting auto-inhibition and stabilizing dynein in an active state. In the presence of Lis1, the occurrence of ∼2.5 pN stalls drops significantly, and DDB-1 complexes predominantly generate ∼4.5 pN forces. Interestingly, higher trap stiffness also promotes this transition, further supporting the role of load in activating dynein^39^. Additionally, D(phi)DB complexes, which are incapable of auto-inhibition^48^, predominantly form two-dynein assemblies (DDB-2) that stall at ∼7 pN, even under sub-molar assembly conditions. This suggests that the open conformation of dynein aligns favorably with an engaged dynein, facilitating the recruitment of additional motors. Interactions between the leading dynein’s LIC and the trailing dynein’s motor domain^69^ (**Fig. 1B**) may further stabilize multi-dynein assemblies. Collectively, our findings underscore how Lis1 and dynein recruitment overcome auto-inhibition to enhance force generation and processivity.

### Tension-Driven Dynein Recruitment

We demonstrate that DDB complexes respond to increasing mechanical load by recruiting a third dynein, raising their stall force to ∼9 pN. This tension-induced recruitment depends on the auxiliary BicD adaptor, which interacts with the LIC of the third dynein. Mutation of the LIC-binding region in the auxiliary BicD(Y46D) significantly reduces third-dynein recruitment, highlighting the importance of this interaction in higher-order DDB assembly.

Structural studies using Cryo-EM^82^ and cryo-ET^41^ have documented DDB assemblies containing one BicD (BICD2) and one dynein, with occasional recruitment of a second dynein under specific conditions^41,42^. Our findings emphasize the pivotal role of mechanical tension in facilitating the recruitment of a third dynein. This enhanced force generation is likely necessary for overcoming cytoplasmic resistance during demanding tasks such as organelle and nuclear transport^83^. Supporting this model, we observed that DDB-2 complexes recruit a third dynein when opposing a MT-bound rigor kinesin, but not in its absence. The measured velocity of ∼30 nm/s for kinesin-DDB-3 complexes matches the velocity of DDB-3 under loads exceeding 8 pN (**Suppl. Fig. 6**). While the precise loading rate at which DDB-3 complexes enhance the force on the kinesin-MT bond remains unclear, the measured velocity suggests that the DDB-3 complex generates sufficient force (∼9 pN) to unbind kinesin from the MT^84^. Furthermore, the constant velocity of the kinesin-DDB-3 complex implies rapid re-binding of kinesin to the MT as the DDB-3 complex continues moving (**Fig. 5C**). These findings strongly support a model in which the DDB complex adapts to increasing loads by recruiting a third dynein—a mechanism consistent with *in vivo* observations of tension-induced dynein recruitment to lipid droplets^85^.

Earlier studies have also shown that Lis1 enhances DDB’s ability to compete with kinesin-1 under load^54^. Without Lis1, only ∼20% of DDB complexes contain two dyneins, resulting in a stall force of ∼4.1 pN, which is insufficient to counter kinesin-1’s stall force of ∼5–6 pN^39,86–88^. Lis1 increases the proportion of two-dynein DDB complexes to ∼40%, raising the stall force to 5.4 pN^54^. Under these conditions, ∼20% of kinesin-DDB complexes exhibit dynein-directed movement, likely due to the recruitment of a third dynein^54^. These findings highlight Lis1’s critical role in enhancing dynein function under load by promoting high-order assembly A recent study on KIF1C-DDB motor complexes reported a 50% increase in DDB processivity when KIF1C was tethered to the complex^89^. Given that KIF1C can generate forces up to ∼7 pN^90^, our observations that DDB-2 complexes stall at ∼7 pN suggests that recruitment of a third dynein, increasing the stall force to ∼9 pN, is required to overcome KIF1C. Together, these results underscore how mechanical tension and Lis1-mediated activation drive the assembly of higher-order dynein complexes, enabling robust force generation and processivity under challenging conditions.

### Dyneins Share Load Unequally Through a Staggered Arrangement

Our results suggest that dyneins within DDB complexes do not share load uniformly. The leading dynein typically slows near its stall force (∼4.5 pN), triggering the engagement of additional motors and rapid increases in force to ∼7 or ∼9 pN. This behavior implies that dyneins are often engaged sequentially as load increases (**Fig. 2G**).

Although cryo-EM tomography shows all four dynein motor domains arranged in parallel^41^, the angled alignment of the dynactin shoulder (∼40° relative to the MT axis) suggests that dyneins bind MTs at offset positions in physiologically relevant conditions^41,42^. Such a “staggered” arrangement would enforce unequal load sharing, consistent with our observed force increments of ∼2.5 pN for additional dyneins. This model is further supported by DNA-origami scaffolds designed to bind two or three DDB-1 complexes in a staggered configuration (with the DNA scaffold aligned to the long MT axis), yielding stall forces of ∼7 pN and ∼9 pN, respectively^60^. Thus, while multiple dyneins contribute to force generation, load sharing among them is inherently unequal.

### DDB Primarily Moves in 8-nm Steps Under Load

We found that DDB complexes, after accounting for their force-dependent compliance (**Suppl. Fig. 7** and **Methods**), predominantly advance with ∼8-nm center-of-mass steps under loads exceeding 2 pN (**Fig. 6B**). This step size remains largely load-independent for DDB-1 and DDB-2, while DDB-3 exhibits smaller forward steps (∼4 nm) as stall forces are approached. These findings suggest a “forward inching” mechanism, where dyneins incrementally shift their positions forward under high loads. This behavior is reminiscent to the stepping behavior of *S. cerevisiae* dynein, which also takes ∼4 nm center-of-mass steps near its stall force^74^. However, the ∼8-nm center-of-mass steps we measured for loads above ∼2 pN are smaller than the previously reported average step size of ∼10 nm under a load of 3.6 pN (step sizes ranging for 2 to 28 nm)^60^. This discrepancy may stem from step-detection algorithms that underfit the data, leading to an overestimation of step sizes while missing smaller increments. By employing an automated step-detection algorithm that eliminates user bias, we aimed to circumvent potential underfitting. Additionally, the use of axonemes rather than MTs in earlier studies may have influenced the stepping behavior of the DDB complexes.

The oscillating forward-backward stepping behavior we identified for human DDB complexes, characterized by a forward step followed by a backward step of relatively short duration, resembles the “transient” backward movements recently reported for *S. cerevisiae* dynein under zero load using MINFLUX microscopy^91^. While the dwell time of DDB complexes after a backward step (∼50 ms) is largely insensitive to loads up to ∼4 pN (**Fig. 6D**), the much shorter dwell time measured in the absence of load (∼4 ms) suggest that the load-induced increase in duration occurs at forces below 0.25 pN (**Fig. 6D**).

It has been shown that the brief backward displacements in the absence of load, which include a diffusional component^91^, correspond to MT release of the leading motor domain. This is followed by a backward “diffusion” of the detached head “around its MT-bound partner” before stepping forward again through a conformational change of its linker^91^. We propose that the detached head undergoes a load-induced backward displacement and, after ATP binding (both events possibly influenced by load), moves forward again through a priming stroke of its linker. Correlative force and single-molecule fluorescence measurements will be needed to confirm whether this is indeed the case.

### Conclusion

In summary, we demonstrate that DDB complexes adapt to increasing mechanical tension by recruiting additional dyneins, thereby boosting force output and processivity. Our findings reveal a novel tension-driven assembly mechanism mediated by the interaction between the auxiliary BicD and dynein’s LIC, linking structural and functional regulation to mechanical adaptation. These insights advance our understanding of intracellular transport and highlight potential therapeutic targets for disorders involving impaired dynein function. Future studies should explore how these mechanisms operate *in vivo* and contribute to cellular homeostasis under diverse mechanical challenges.

## Materials and Methods

### Plasmid and Construct Generation

The plasmids for human dynein complexes were generously provided by Dr. Andrew P. Carter (MRC Laboratory of Molecular Biology, Cambridge, UK). Plasmids for Lis1 constructs were gifts from Dr. Samara Reck-Peterson (University of California, San Diego, CA). BicD2 constructs were generated using a mouse BicD2 plasmid (Addgene, #64205)^92^ and either a backbone optimized for *E. coli* expression (pSNAP-tag(T7)-2 vector, New England Biolabs, #N9181S)^86^ or pET28b. Shorter BicD2 constructs were produced via Q5 mutagenesis (New England Biolabs, # E0554S) and a single mutation (Y46D) was introduced into the BicD2(25-180) construct.

The kinesin (Cys-light) S43C construct^93^ was kindly provided by Dr. Ronald D. Vale (previously University of California, San Francisco, CA). To generate the rigor kinesin construct, a single mutation (T92N) was introduced, and a SpyTag003 was inserted at the C-terminus using Q5 mutagenesis. The SpyCatcher003 tag, amplified from the SpyCatcher003-S49C plasmid^73^ (Addgene, #133448), was inserted to the C-terminus of BicD2. Plasmids for Tev protease^94^ (Addgene, #171782) and BirA (Addgene, #20857)^95^ were obtained from Addgene.

All plasmids generated for this study were validated through Sanger sequencing (Genomics Core Facility, Albert Einstein College of Medicine, Bronx, NY) or full-plasmid sequencing (Azenta). The constructs used in this study are listed in Supplementary Table 1.

### Tagging Dynactin in HEK 293T/17 SF Cells Using CRISPR

A HaloTag-3×FLAG-P2A-mCherry tag was inserted at the C-terminus of the DCTN4 (p62) subunit of dynactin using a CRISPR-based method adapted from a previously published protocol^65^. The HaloTag-3×FLAG has been used successfully in a stable adherent HEK293 cell line for dynactin purification^55^. The DCTN4 gene and 1 kb downstream sequence were retrieved from the UCSC Genome Browser (https://genome.ucsc.edu/), and the guide RNA was designed using the CRISPOR online tool (https://crispor.gi.ucsc.edu/)^96^. The guide RNA (5’-TCCTTAAAAGGTTCCACTGG) was ordered as Alt-R® crRNA from IDTDNA, along with Alt-R® tracrRNA (IDTDNA, #1072532) and Alt-R® Cas9 nuclease (IDTDNA, #1081058). These RNAs were dissolved in nuclease-free duplex buffer (IDTDNA) to final 100 µM.

To construct the targeting plasmid, 303 bp upstream of the DCTN4 stop codon were amplified from HEK 293T/17 SF genomic DNA and stitched with HaloTag-P2A-mCherry, followed by 611 bp of downstream DCTN4 sequences. The resulting DNA fragment was then inserted into a backbone plasmid, and the construct was confirmed by full plasmid sequencing (Azenta).

### Cell Culture and Preparation

One milliliter of HEK 293T/17 SF cells (ATCC, #ACS-4500) was recovered following the vendor’s instructions in 10 mL of complete medium (9.8 mL of BalanCD HEK293 (Fujifilm Irvine Scientific, #91165-1L), 0.1 mL of 100×GlutaMAX™ supplement (Gibco, #35050061), 0.1 mL of 100×ITS (Insulin-Transferrin-Selenium, Corning, #25-800-CR), and 20 µL of anti-clumping agent (Gibco, #0010057DG)). Cultures were shaken at 37°C, 125 rpm, with 5% CO_2_ (Eppendorf, New Brunswick S41i CO2 Incubator Shaker) and maintained at a density of < 2×10^6^ cells/mL.

### Electroporation and CRISPR Delivery

For CRISPR delivery, 3 µL of crRNA, 3 µL of tracrRNA, and 4 µL of Neon™ NxT resuspension R buffer (ThermoFisher, # NEON1S) were mixed, annealed by being heating at 95°C for 5 minutes, and then cooled to room temperature (RT). Cas9 protein (1.5 µL of the 10 mg/mL) was added to the RNA mixture and incubated at RT for 15 minutes, followed by the addition of 6 µg of the targeting plasmid.

HEK 293T/17 SF cells (5×10^6^) were pelleted at 140×g for 5 minutes, washed with 10 mL PBS, and then resuspended in 100 µL of Neon™ NxT R Buffer. The prepared Cas9-RNA-plasmid complex was mixed with the resuspended cells, and electroporation at 1100 V for 20 ms with 2 pulses using the Neon™ NxT Electroporation System (ThermoFisher, # NEON1S), following the manufacturer’s instruction. The electroporated cells were immediately transferred to 2 mL of complete medium in a 6-well plate and allowed to recover for 2 days at 37°C without shaking. After recovering, approximately 10^7^ cells were sorted using a BD Aria cell sorter to select for mCherry-positive cells. The sorted cells were then cultured, frozen, and stored in liquid nitrogen for future use.

### Dynactin Purification

HEK 293T/17 SF cells expressing p62 tagged with HaloTag-3×FLAG were cultured in 800 mL of medium at a density of 2×10^6^ cells/mL and harvested by centrifugation at 600×g for 10 minutes at 4°C. The resulting pellet (∼10 mL) was resuspended in 10 mL of 2×dynactin lysis buffer (DLB) containing 60 mM HEPES (pH 7.2), 200 mM NaCl, 4 mM MgCl_2_, 2 mM EGTA, 20% glycerol, 0.2 mM ATP, 2 mM DTT, 0.4% (v/v) Triton X-100, and two EDTA-free protease inhibitor cocktail tablets (Roche, #11836170001). The suspension was nutated at 4°C for 15 minutes. The lysate was then cleared by centrifugation at 80,000 rpm (260,000×g, *k*-factor=28) for 10 minutes using a TLA-110 rotor in a Beckman Tabletop Optima TLX Ultracentrifuge.

Anti-FLAG® M2 affinity gel (1 mL; Sigma, #A2220) was washed with 5 mL of 1×DLB buffer. The cleared lysate was added to the resin and nutated overnight at 4°C. The resin was subsequently washed with 40 mL of wash buffer containing 30 mM HEPES (pH 7.2), 250 mM KCl, 2 mM MgCl_2_, 1 mM EGTA, 10% glycerol, 0.1 mM ATP, 1 mM DTT, 0.2% (w/v) Pluronic F-127, and 0.5 mM Pefabloc. The resin was then transferred to a 2-mL tube and incubated with 200 µL of wash buffer containing 100 µL of 5 mg/mL FLAG peptide. This mixture was nutated at 4°C for 30 minutes. The supernatant containing the purified dynactin was collected and concentrated using an Amicon Ultra-0.5 mL centrifugal filter unit (100-kDa MWCO) at 5000×g at 4°C. The concentration of dynactin was determined using a Bradford assay (ThermoFisher, #23200). The subunit composition of dynactin was confirmed by mass spectrometry.

### Dynactin From Porcine Brain

Porcine brain dynactin was kindly provided by Dr. Andrew P. Carter (MRC, Laboratory of Molecular Biology, Cambridge, UK).

### Sf9 Expression of Cytoplasmic Dynein-1 and Lis1

The full cytoplasmic dynein-1 complex, the tail complex, the phi-mutant complex, and Lis1 were expressed in Sf9 cells following established protocols^25,55^. Briefly, a bacmid was generated by transforming the plasmid containing the desired construct into MAX Efficiency™ DH10Bac competent cells (Gibco, # 10361012). Recombinant bacmid insertion was verified by blue-white screening and confirmed by PCR.

One milliliter of Sf9 cells cultured in Sf-900™ II SFM (ThermoFisher, #11496015) was recovered and grown in 10 mL of Sf-900™ II SFM medium (ThermoFisher, #10902104) at 27°C with shaking at 135 rpm. Cultures were maintained at a density of < 2×10^6^ cells/mL. For transfection, 2 mL of a 0.5×10^6^ cells/mL culture was seeded in a 6-well plate. A mixture of 2 µg of bacmid and 200 µL of medium was prepared, and 6 µL of FuGENE® HD transfection reagent (Promega, #E2311) was added.

After a 15-minute incubation at RT, the mixture was added dropwise to the cells. The culture was then incubated at 27°C without shaking for 4 days to generate P1 virus. The P1 virus was collected, transferred to a 15-mL tube, and stored at 4°C in the dark. To produce P2 virus, 0.5 mL of the P1 virus was added to 50 mL of 1.5×10^6^ cells/mL culture and incubated at 27°C with shaking at 135 rpm for 3 days. The P2 virus was harvested by centrifugation at 2000×g for 10 minutes at 4°C and stored at 4°C in the dark. For protein expression, 5 mL of the P2 virus was added to 500 mL culture of Sf9 cells at a density of 2×10^6^ cells/mL. The culture was shaken at 27°C at 135 rpm for 60–72 hours. Cells were harvested by centrifugation at 2000×g for 15 minutes at 4°C. The pellet was washed with 40 mL of cold PBS and centrifuged again at 2000×g for 10 minutes at 4°C. The supernatant was discarded, and the pellet was flash frozen in liquid nitrogen and stored at –80°C until further use.

### Cytoplasmic Dynein-1 and Lis1 Purification

The cell pellet (∼5 mL) was resuspended in 5 mL of 2×lysis buffer containing 60 mM HEPES (pH 7.2), 300 mM KCl, 4 mM MgCl_2_, 2 mM EGTA, 20% (v/v) glycerol, 0.4 mM ATP, 4 mM DTT, 0.2% (w/v) Pluronic F-127, 100 µL of 2.5 U/µL Dnase I, and two EDTA-free protease inhibitor cocktail tablets. The resuspended pellet was dounced for 30 strokes on ice. The lysate was cleared by centrifugation as described for dynactin purification. Four milliliters of IgG Sepharose 6 Fast Flow affinity resin (Cytiva, #17096901) was pre-washed with 6 mL of lysis buffer. The cleared lysate was added to the resin and nutate at 4°C for 4 hours. The resin was then washed with 50 mL of wash buffer containing 50 mM HEPES (pH 7.2), 150 mM KCl, 2 mM MgCl_2_, 1 mM EGTA, 10% (v/v) glycerol, 0.1 mM ATP, 1 mM DTT, 0.1% (w/v) Pluronic F-127, and 1 mM PMSF.

To label the SNAP-tag on the N-terminus of the dynein heavy chain, 15 µL of 1 mM SNAP-tag dye was added to the resin, and the mixture was nutated at 4°C overnight. The resin was then washed with 50 mL of TEV-release buffer containing 50 mM HEPES (pH 7.2), 150 mM KCl, 2 mM MgCl_2_, 1 mM EGTA, 10% (v/v) glycerol, 0.1 mM ATP, 1 mM DTT, and 0.1% (w/v) Pluronic F-127. The resin was subsequently transferred to a 5-mL tube with TEV-release buffer to a final volume of 5 mL. Fifty microliters of 200 µM MBP-superTevProtease was added to the resin, and the mixture was nutated at 4°C overnight in the dark. One milliliter of amylose resin (New England Biolabs, #E8021S) was used to remove the MBP-superTev protease. The supernatant was concentrated following the same protocol used for dynactin purification. The purity of the cytoplasmic dynein-1 complex and Lis1 was verified by 4-12% bis-tris polyacrylamide gel electrophoresis, and the concentration was determined using the Bradford assay.

### *E. coli*-Based Expression

BicD2 constructs were expressed in *E. coli*. Each plasmid was transformed into BL21-CodonPlus(DE3)-RIPL competent cells (Agilent Technologies, #230280), and a single colony was picked and inoculated in 1 mL of terrific broth (TB) containing 50 µg/mL carbenicillin and 50 µg/mL chloramphenicol. The 1-mL culture was shaken at 37°C overnight and subsequently inoculated into 400 mL of TB supplemented with 2 µg/mL carbenicillin and 2 µg/mL chloramphenicol. The culture was shaken at 37°C for 5 hours and cooled to 16°C for 1 hour. Protein expression was induced with 0.1 mM IPTG continued overnight at 16°C. The cells were harvested by centrifugation at 3,000×g for 10 minutes at 4°C, and the supernatant was discarded. The cell pellet was resuspended in 5 mL of B-PER™ Complete Bacterial Protein Extraction Reagent (ThermoFisher Scientific, #89821) supplemented with 2 mM MgCl_2_, 1 mM EGTA, 1 mM DTT, 0.1 mM ATP, and 2 mM PMSF. The resuspension was flash-frozen in liquid nitrogen and stored at –80 °C.

### Purification of *E. coli*-Expressed Constructs

To purify *E. coli*-expressed protein, the frozen cell pellet was thawed at 37 °C and nutated at RT for 20 minutes to lyse the cells. The cell lysate was cleared as for dynactin purification. The supernatant was passed through 500 μL of Ni-NTA Roche cOmplete™ His-Tag purification resin (Millipore Sigma, #5893682001) for His-tag tagged proteins or Strep-Tactin® 4Flow® high-capacity resin (IBA Lifesciences GmbH, #2-1250-010) for strep-II tagged proteins. The resin was washed with 10 mL of wash buffer containing 50 mM HEPES (pH 7.2), 300 mM KCl, 2 mM MgCl_2_, 1 mM EGTA, 1 mM DTT, 1 mM PMSF, 0.1 mM ATP, 0.1% (w/v) Pluronic F-127, and 10% glycerol. Proteins were eluted with elution buffer (50 mM HEPES, pH7.2, 150 mM KCl, 2 mM MgCl_2_, 1 mM EGTA, 1 mM DTT, 1 mM PMSF, 0.1 mM ATP, 0.1% Pluronic F-127 (w/v), 10% glycerol) supplemented with either 150 mM imidazole for His-tagged proteins or 5 mM desthiobiotin for strep-II tagged proteins. The elute was either flash frozen and stored at –80 °C or concentrated using an Amicon Ultra-0.5 mL centrifugal filter unit (30-kDa MWCO) (Sigma, #UFC503024) before flash freezing. The purity and concentration of the proteins were analyzed using 4-12% bis-tris polyacrylamide gels. For biotinylation of BicD2, 50 µL of 10 µM BicD2 was mixed with 1 µL of 100 mM ATP, 2 µL of 50 µM BirA, and 1 µL of 10 mM biotin, and incubated at 30°C for 2 hours. BirA was subsequently removed using Ni-NTA resin and the solution was buffer exchanged into 30 mM HEPES (pH 7.2), 50 mM KCl, 2 mM MgCl_2_, 1 mM EGTA, 1 mM DTT, 0.1% (w/v) Pluronic F-127, and 10% glycerol to remove free biotin.

### Purification of the BicD2-Kinesin Constructs

The purification of BicD2-kinesin constructs followed the same protocol as for *E. coli*-expressed constructs, with modifications to facilitate complex formation. After the cells were thawed, the BicD2 and kinesin solutions were combined and nutated at RT for 20 minutes, followed by additional nutation at 4°C for 2 hours to allow binding between SpyCatcher003 and SpyTag003. The lysate was then passed through Ni-NTA resin to capture His-tagged components. The elution from the Ni-NTA resin was subsequently passed through Strep-Tactin resin to isolate the fully formed BicD2-kinesin complex. The final elute from the Strep-Tactin resin was further concentrated following the protocol used for dynactin purification.

### Optical Tweezers Assay

#### Polystyrene Beads

Anti-GFP polystyrene beads were prepared by covalently binding anti-GFP antibodies and BSA to carboxyl polystyrene beads (0.52 µm, Polysciences, #09836-15) using NHS-EDAC chemistry^80^. Streptavidin beads (0.55 µm) were purchased from Spherotech (#SVP-05-10).

#### Microtubule Polymerization

Two microliters of 10 mg/mL tubulin (Cytoskeleton, #T240-B) were mixed with 2 µL of 1 mg/mL biotinylated tubulin (Cytoskeleton, # T333P-A) and 1 µL of 10 mM GTP. The mixture was incubated at 37°C for 20 minutes. After incubation, 0.5 µL of 0.2 mM paclitaxel in DMSO was added, and the incubation was continued for an additional 20 minutes. The solution was carefully layered on top of a 100 µL glycerol cushion (80 mM PIPES, pH 6.8, 2 mM MgCl_2_, 1 mM EGTA, 60% (v/v) glycerol, 1 mM DTT, and 10 µM paclitaxel) in a 230-µL TLA100 tube (Beckman Coulter, #343775) and centrifuged at 80,000×rpm (250,000×g, *k*-factor=10) for 5 minutes at RT. The supernatant was gently removed, and the pellet was resuspended in 11 µL of BRB80G10 (80 mM PIPES, pH 6.8, 2 mM MgCl_2_, 1 mM EGTA, 10% (v/v) glycerol, 1 mM DTT, and 10 µM paclitaxel). The MT solution was stored at RT in the dark for further use.

#### Flow Chamber Preparation

A flow chamber was assembled using a glass slide (Fisher Scientific, #12-550-123) and an ethanol-cleaned coverslip (Zeiss, #474030-9000-000), separated by two thin strips of parafilm. Ten microliters of 0.5 mg/ml BSA-biotin (ThermoScientific, #29130) were introduced into the chamber and incubated for 10 minutes. The chamber was then washed with 2×20 µL blocking buffer (80 mM PIPES, pH 6.8, 2 mM MgCl_2_, 1 mM EGTA, 10 µM paclitaxel, 1% (w/v) Pluronic F-127, 2 mg/mL BSA, and 1 mg/mL α-casein) and incubated for 30 minutes to block the surface. Next, 10 µL of 0.25 mg/ml streptavidin (Promega, #Z7041) was introduced into the chamber and incubated for 10 minutes. The chamber was washed with 2×20 µL blocking buffer, followed by the addition of 10 µL of 0.02 mg/ml biotin-labeled MTs in the blocking buffer, which were incubated for 1 minute. After another wash with 2×20 µL blocking buffer, the chamber was stored in a humidity chamber until further use.

#### Sample Preparation

Dynein (dimer), dynactin, and BicD2 (dimer) were mixed in a 1:1:1 ratio to a final concentration of 200 nM each. The solution was incubated at 4°C overnight to allow complex formation. One microliter of polystyrene trapping beads was mixed with 1 µL of motility buffer (60 mM HEPES, pH 7.2, 50 mM KCl, 2 mM MgCl_2_, 1 mM EGTA, 10 µM paclitaxel, 0.5% (w/v) Pluronic F-127, 5 mg/mL BSA, and 0.2 mg/mL α-casein) and 1 µL of appropriately diluted DDB complex. The mixture was incubated on ice for 30 minutes. Following incubation, 40 µL of motility buffer supplemented with 2 mM ATP, 2 mM biotin, and a gloxy oxygen scavenger system was added to the bead-protein solution. Two 20 µL volumes of the solution were then introduced into the chamber. The chamber was sealed with vacuum grease to prevent evaporation.

#### Data Acquisition

Optical tweezers experiments were performed using a C-Trap combined with total internal reflection fluorescence (TIRF) and interference reflection microscopy (IRM) (C-Trap Edge, LUMICKS) at RT. MTs were visualized using IRM. Trap stiffness was adjusted to ∼0.08–0.1 pN/nm. Optical trapping data were sampled at a rate of 20 MHz and digitally down-sampled by a factor of 256, resulting in a final sample rate of 78.125 kHz. An anti-aliasing filter was applied to achieve a passband of 31.25 kHz. The resulting data were fitted to a Lorentzian power spectrum for analysis.

#### Data Analysis

Data were processed using a custom MATLAB program. A built-in MATLAB step-finding algorithm was applied to traces that were down-sampled by a factor of 80 through averaging. Steps with forces below 0.2 pN were excluded. For each step, the combined stiffness of the motor and bead-linkage, *k_m-link_*, was calculated using the original data^97^:

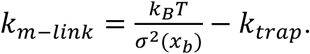

Steps for each trace were binned into 0.5-pN force windows. The mean value of the 25^th^ to 75^th^ percentiles for each force window (e.g., dwell time, step sizes and velocity) were calculated to represent in the force window. Traces with similar forces were grouped, and the mean and standard error of the mean (SEM) were calculated and visualized using Prism 10 (GraphPad). To compensate for the linkage compliance, the difference between the motor stretch (F/*k_m-link_*) for each force window in each group was calculated and added to the steps.

### Total Internal Reflection Fluorescence (TIRF) Assay

#### Sample Preparation

MT polymerization and flow chamber preparation were performed as described for the optical tweezers assay. After MTs were immobilized on the cover-slip surface, the assembled complex was diluted in the motility buffer containing 60 mM HEPES (pH 7.2), 75 mM KCl, 2 mM MgCl_2_, 1 mM EGTA, 1 mM DTT, 2 mM ATP, 0.5% Pluronic F-127 (w/v), and 10% glycerol. The solution was introduced into the slide chamber, which was then sealed with vacuum grease to prevent evaporation.

For three-color TIRF experiments, the DDB complexes were pre-assembled overnight in in motility buffer containing 100 µM ATP, with two dyneins labeled with different dyes (Alexa Fluor 647 and TMR dyes). After constructing of the slide chamber and immobilizing MTs, 10 nM dynein labeled with Alexa Fluor 488 was added to the pre-assembled DDB complexes and flown into the chamber.

#### Data Acquisition

Images were acquired using BioVis software (BioVision Technologies) with an acquisition time of either 200 ms or 500 ms per frame.

#### Data Analysis

Kymographs were generated using Fiji^98^, and velocities were analyzed using a custom MATLAB program.

### AlphaFold2 Structure Prediction

The interaction between mouse BicD2(25-180) and human LIC2(417-444) was predicted using ColabFold (v1.5.5: AlphaFold2 using MMseqs2)^99^.

### Graphic Visualization

Molecular structures in the figures were visualized using UCSF ChimeraX^100^. Model graphics in Figures 2 and 7 were generated using BioRender (https://www.biorender.com/).

**Figure 7.**
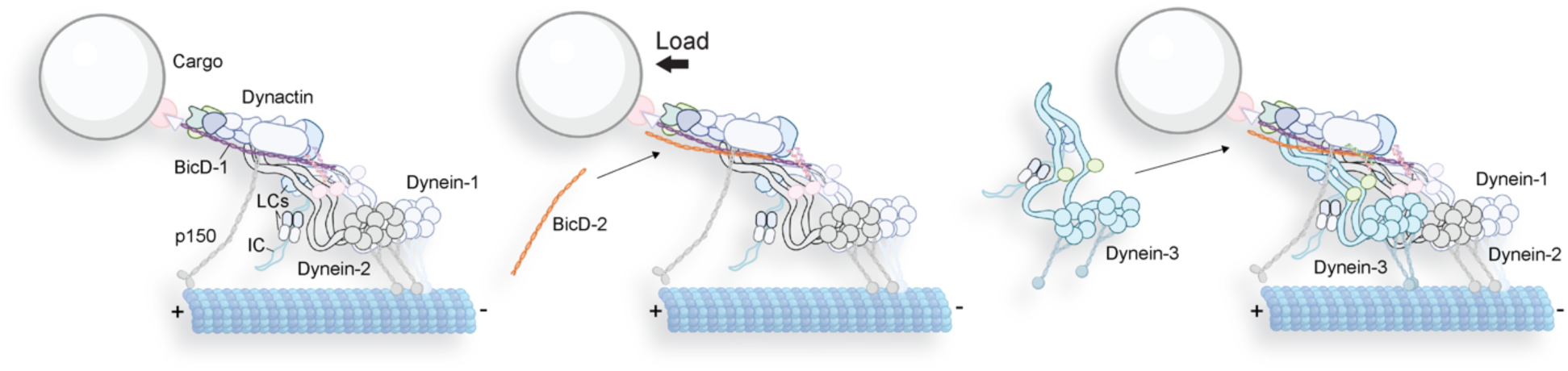
Proposed model of a third dynein dimer being recruited to a DDB complex through a second BicD molecule under applied force.

## Acknowledgments

We acknowledge Albert Einstein College of Medicine Flow Cytometry Core Facility (Einstein National Cancer Institute’s Cancer Center support grant P30CA013330), especially Ming Liu, for assistance with sorting HEK293 cells. We also thank Jan O. Wirth (Abberior) for valuable discussions on step-finding algorithms. L. Rao and A. Gennerich were supported by National Institutes of Health grants R01GM098469 and would like to acknowledge the use of LUMICKS C-Trap, funded by the NIH grant S10OD034445-01. R. McKenney was supported by NIGMS National Institutes of Health grant GM124889. M. Arnold and K. Stengel were supported by National Institutes of Health grant R35GM147213. S. Sidoli gratefully acknowledges the funding from the Hevolution Foundation (AFAR), the Einstein-Mount Sinai Diabetes Center, and the NIH Office of the Director (S10OD030286). Molecular graphics and analyses performed with UCSF ChimeraX, developed at the University of California, San Francisco, with support from NIH R01-GM129325 and the Office of Cyber Infrastructure and Computational Biology, National Institute of Allergy and Infectious Diseases.

## Author Contributions

L. Rao produced and purified all proteins; L. Rao and A. Gennerich designed the research; L. Rao and X. Liu expressed and purified proteins; L. Rao, F. Berger, and A. Gennerich analyzed the experimental data; L. Rao and A. Gennerich wrote the manuscript. A. Gennerich secured funding; R. McKenney generated and provided proteins purified from rat brain lysate for initial studies. M. Arnold and K. Stengel assisted in CRISPR-based dynactin construct generation; S. Sidoli performed the mass spectrometry assay for dynactin.

## Data Availability

Data supporting the findings of this manuscript are available from the corresponding author upon reasonable request.

