## Supplementary Figures and Table for "The Power of Three: Dynactin associates with three dyneins under load for greater force production"

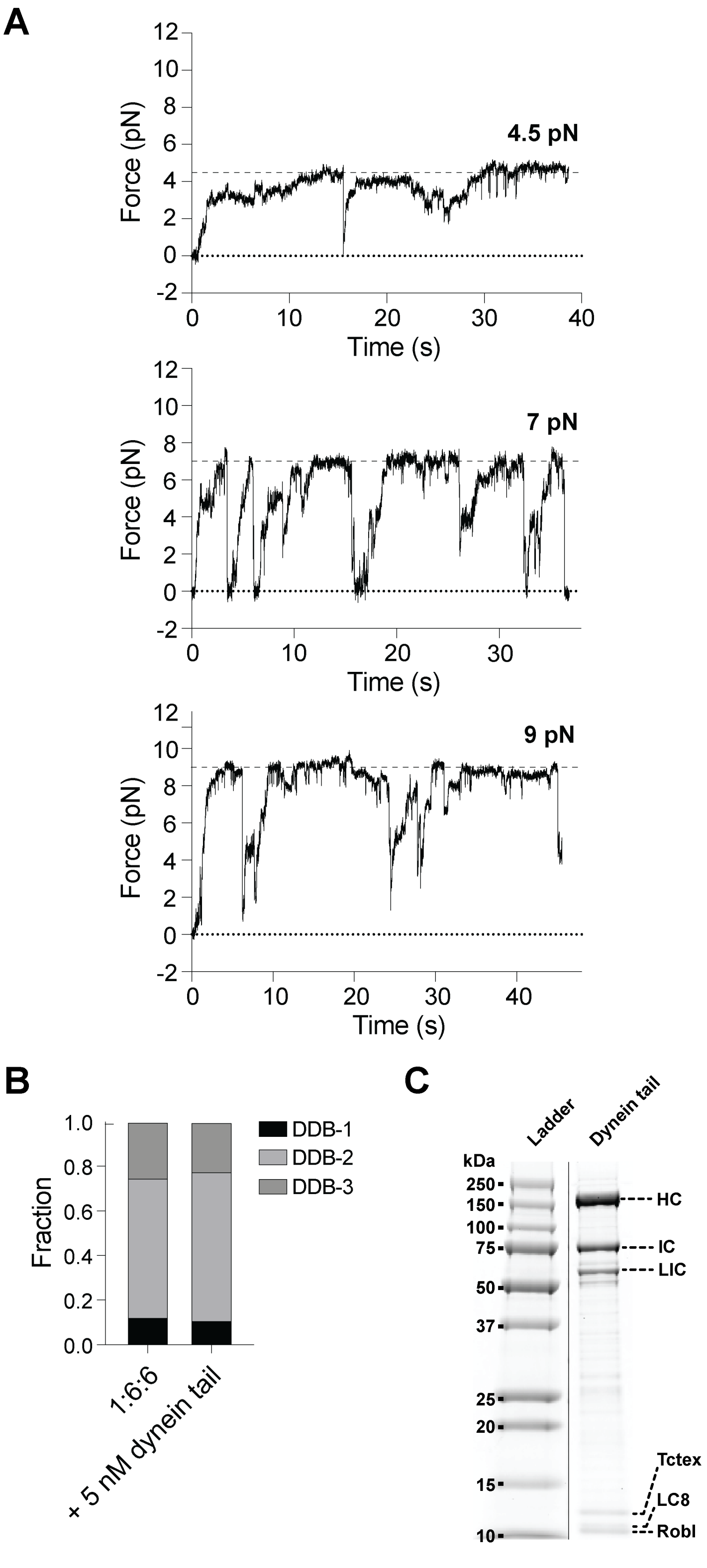

**Supplemental Figure 1. (A)** An example traces of D(phi)DB generating ~4.5 pN, ~7 pN and ~9 pN force. The traces have been down-sampled by a factor of 400 for representation. **(B)** The fraction of beads showing different force generation behavior. The percentages represent DDB-1 (fully activated), DDB-2, and DDB-3, respectively. 1:6:6 (dynein(phi):dynactin:BicD): n= 9, 12%, 62%, 26%. +5 nM dynein tail: n=8, 11%, 67%, 22%. (**C**) Gel showing the purified dynein tail.

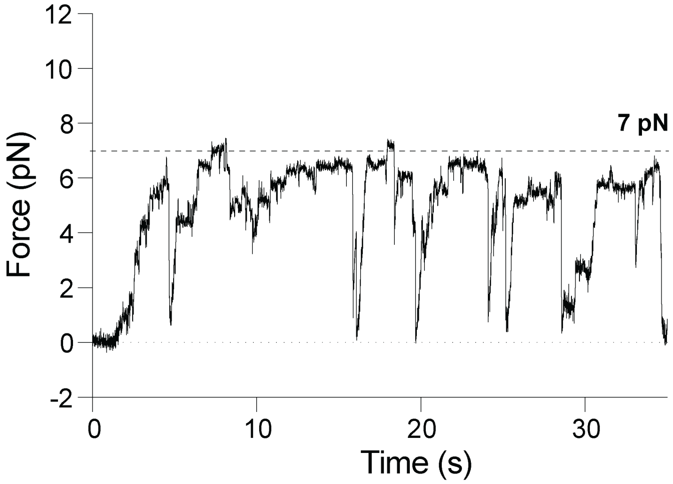


**Supplemental Figure 2.** An example trace of DDB generating ~7 pN force in the presence of 5 nM free dynein. The trace has been down-sampled by a factor of 400 for representation.

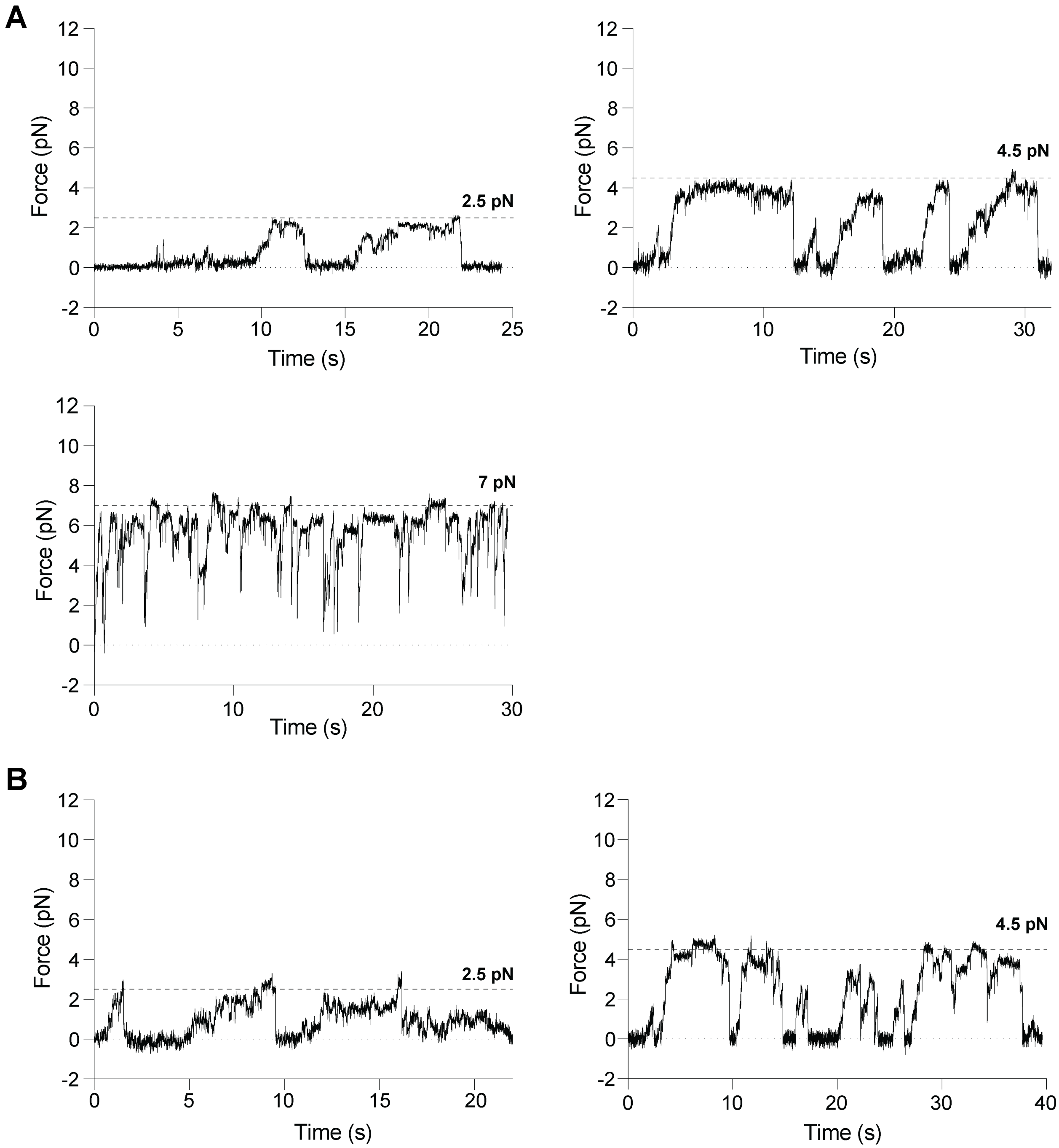


**Supplemental Figure 3. (A)** An example traces of DDB(276) generating ~2.5 pN, ~4.5 pN, and ~7 pN force. **(B)** An example trace of DDB(180) generating ~2.5 pN and ~4.5 pN force. The traces have been down-sampled by a factor of 400 for representation.

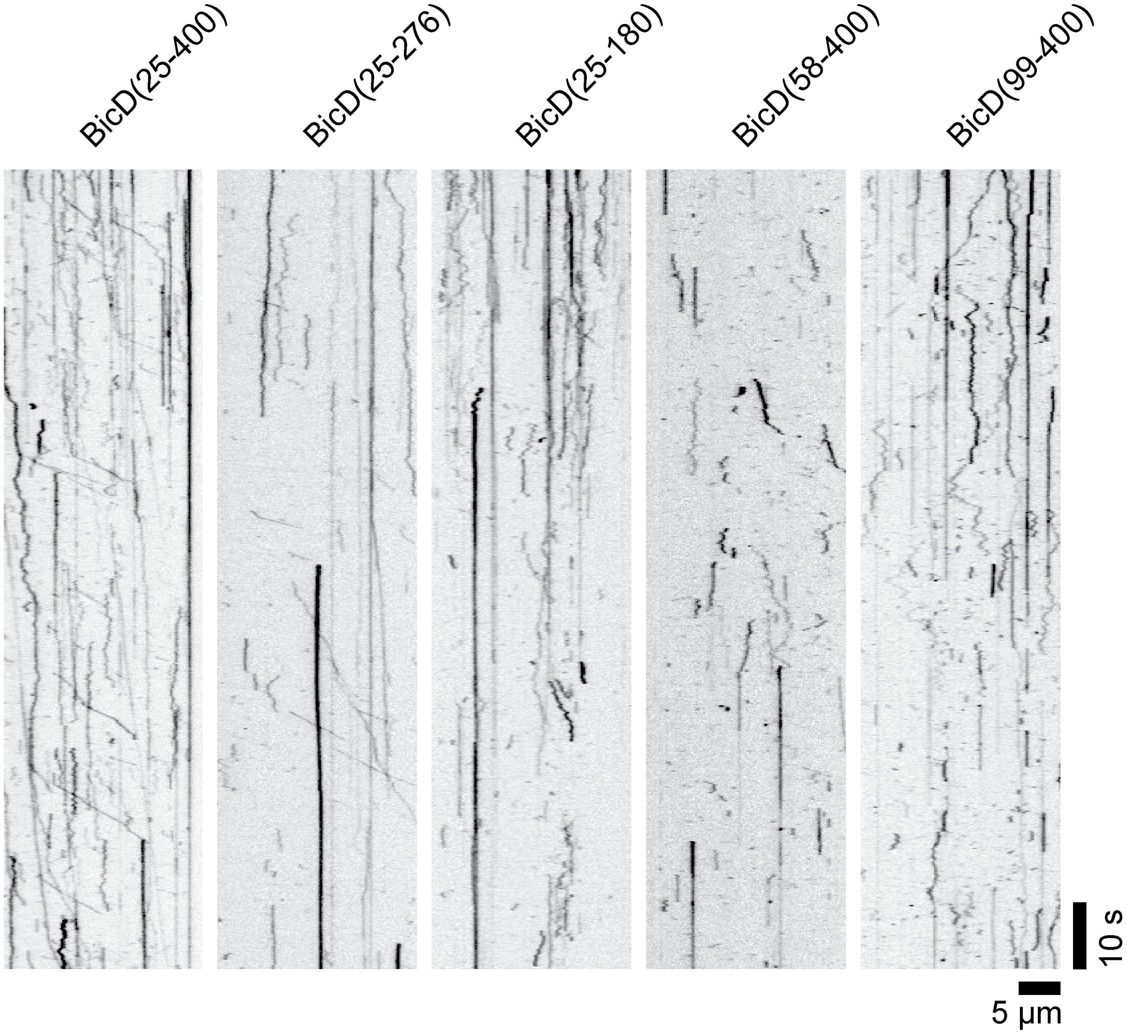


**Supplemental Figure 4.** Kymograph examples of the motility of DDB complexes assembled with different lengths of BicD.

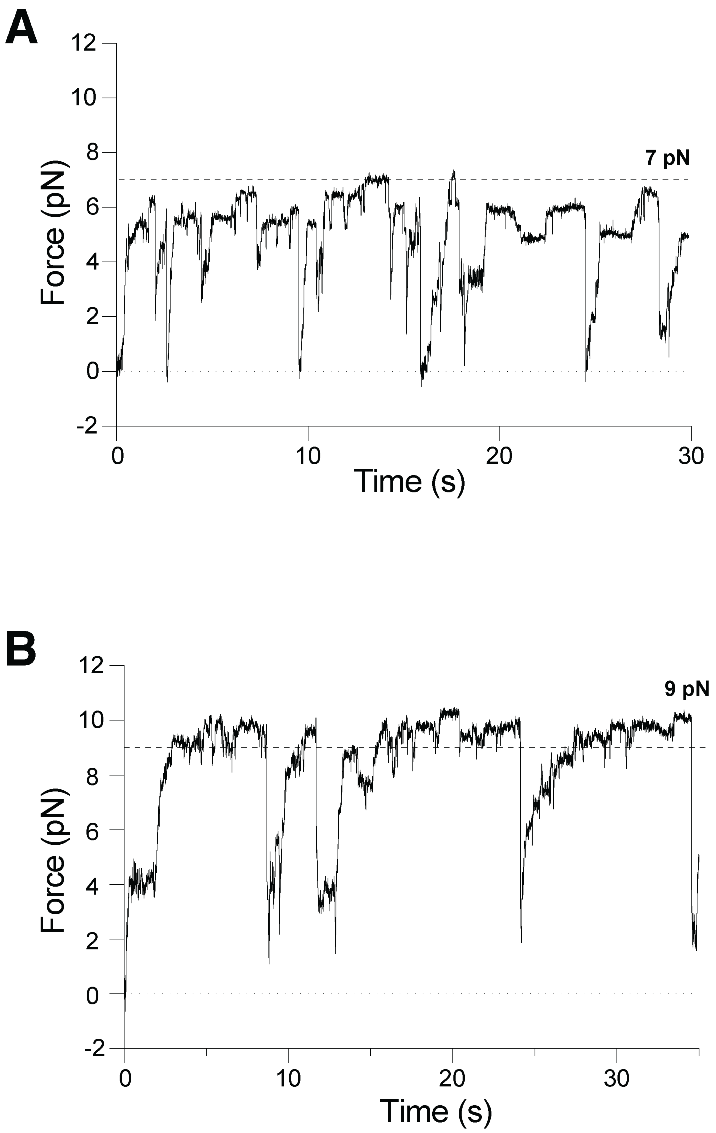


**Supplemental Figure 5. (A)** An example trace of DDB(400) generating ~7 pN force in the presence of 5 nM dynein and 5 nM BicD(180). **(B)** An example trace of DDB(400) generating ~9 pN force in the presence of 5 nM dynein and 5 nM BicD(180). The traces have been down-sampled by a factor of 400 for representation.

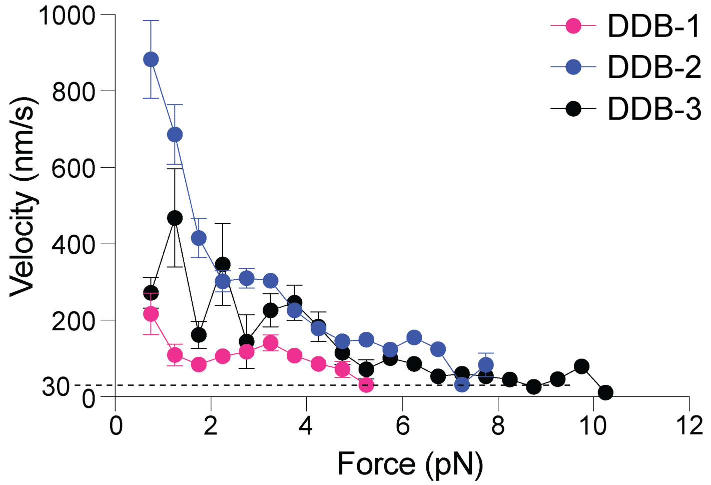


**Supplemental Figure 6.** Velocity of DDB complexes as a function of force. The bars represent mean with SEMs. Red: DDB-1; blue: DDB-2; black: DDB-3. The velocity was derived from compliance-corrected step sizes and dwell times. A window of 0.5-pN force was used to bin the data.

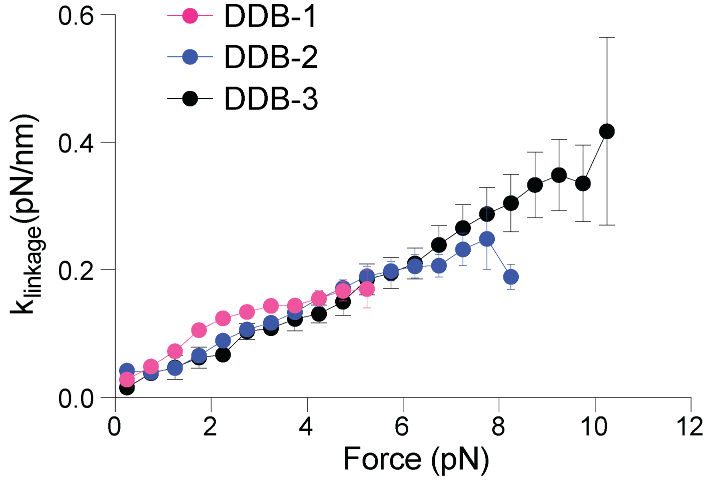


**Supplemental Figure 7.** Measured $k_{linkage}$ of DDB complexes as a function of force. Bars represent mean values, with error bars indicating the SEM. Red: DDB-1; blue: DDB-2; black: DDB-3.

**Supplemental Table 1.** Constructs used in this study.

| Protein | Construct | Expression System | |
|---|---|---|---|
| Human cytoplasmic dynein-1 full complex | His-ZZ-Tev-SNAPf DHC1_IC2C_LIC2_Tctex1_Robl1_LC8 | Sf9 | 10.15252/embj.201488792 |
| Human cytoplasmic dynein-1 tail | His-ZZ-Tev-SNAPf DHC1(1-1074)_IC2C_LIC2_Tctex1_Robl1_LC8 | Sf9 | 10.1038/nature25462 |
| Human cytoplasmic dynein-1 phi-mutant | His-ZZ-Tev-SNAPf DHC1(K1610E, R1567E)_IC2C_LIC2_Tctex1_Robl1_LC8 | Sf9 | 10.1016/j.cell.2017.05.025 |
| Human Lis1 | ZZ-Tev-Lis1 | Sf9 | 10.1038/s41556-020-0506-z |
| Human dynactin | p62-HaloTag-3×FLAG | HEK293T | This study |
| Mouse BicD2 | BicD2(25-400)-sfGFP-StrepII | *E. coli* | This study |
| Mouse BicD2 | BicD2(25-400)-AviTag-StrepII | *E. coli* | This study |
| Mouse BicD2 | BicD2(58-400)-AviTag-StrepII | *E. coli* | This study |
| Mouse BicD2 | BicD2(99-400)-AviTag-StrepII | *E. coli* | This study |
| Mouse BicD2 | BicD2(25-276)-AviTag-StrepII | *E. coli* | This study |
| Mouse BicD2 | BicD2(25-180)-HaloTag-StrepII | *E. coli* | This study |
| Mouse BicD2 | BicD2(25-180, Y46D)-HaloTag-StrepII | *E. coli* | This study |
| Mouse BicD2 | BicD2(25-400)-SpyCatcher003-StrepII | *E. coli* | This study |
| Human kif5b | kif5b(490)(cys-light, S43C, T72N)-SpyTag003-6His | *E. coli* | This study |
